# CoRegNet: Unraveling Gene Co-regulation Networks from Public RNA-Seq Repositories Using a Beta-Binomial Statistical Model

**DOI:** 10.1101/2022.10.17.512527

**Authors:** Jiasheng Wang, Ying-Wooi Wan, Rami Al-Ouran, Meichen Huang, Zhandong Liu

## Abstract

Millions of RNA sequencing samples have been deposited into public databases, providing a rich resource for biological research. These datasets encompass tens of thousands of experiments and offer comprehensive insights into human cellular regulation. However, a major challenge is how to integrate these experiments that acquired at different conditions. We propose a new statistical tool based on beta-binomial distributions that can construct robust gene co-regulation network (CoRegNet) across tens of thousands of experiments. Our analysis of over 12,000 experiments involving human tissues and cells shows that CoRegNet significantly outperforms existing gene co-expression-based methods. Although the majority of the genes are linearly co-regulated, we did discover an interesting set of genes that are non-linearly co-regulated; half of the time they change in the same direction and the other half they change in the opposite direction. Additionally, we identified a set of gene pairs that follows the Simpson’s paradox. By utilizing public domain data, CoRegNet offers a powerful approach for identifying functionally related gene pairs, thereby revealing new biological insights.

## INTRODUCTION

Since the development of high throughput sequencing technology, tens of thousands of experiments have used RNA sequencing technology to measure gene expression changes in response to genetic, chemical and environmental regulation. Majority of these samples are now available in the public data repositories such as GEO and ArrayExpress[1,2]. Many bioinformatics tools have been developed to query the gene expression changes in each individual experiment. However, little has been done to integrate these data together due to the vast heterogeneous nature of the experiments.

Correlation based bioinformatics tool such as the widely used Weighted Gene Co-expression Network Analysis (WGCNA) can identify co-regulated genes among a large number of samples; However, these samples need to be acquired under similar conditions or adjusted using batch correction algorithms.

Although tools like limma and ComBat-seq can correct batch effects[3,4], the batch factor is completely confounded with the experimental designs making these approaches not applicable. Models like mixed model co-expression (MMC) can identify co-expression patterns even in heterogeneous expression data, they over-correct true biological signals by interpreting hub genes’ biological effect as a global confounder and adjusting it[5]. Mutual Information (MI) serves as a robust, nonparametric statistical methodology proficient in delineating gene-gene associations[6]. However, its efficacy diminishes when confronted with heterogeneous co-expression patterns across different experimental conditions [Sup Figure 1].

To circumvent the batch issue, several studies have tried to combine samples from a small number of studies (20-100 studies) that done in similar conditions, which may help to reduce the batch effect’s impact on results[7–10]. However, it is an ill-posed problem to compute Pearson’s correlation over experimental designs involves two groups of treatment and controls. Samples in the same group tend to cluster together and may even overlap with each other, resulting in an inflated statistical significance when computing correlation values [Sup Figure 2]. In addition, the correlations between different treatment and control groups differ widely, so it is difficult to combine them together. The widely used co-expression model is therefore inapplicable to identify gene-gene associations [Sup Animation 1].

Accurate estimation of Pearson’s correlation between a pair of genes requires a large number of samples under similar experimental conditions. In Recount3, 6,377 out of 8,677 SRA human studies have less than 20 samples. In other words, if a study has two groups, each group typically has ≤ 10 samples— if it contains multiple groups, this number is even smaller. Such small sample sizes make it difficult to estimate gene correlations in a single study[11].

Here, we propose a new co-regulation model -CoRegNet. This model generates a gene co-regulation network in which genes co-regulated under many heterogeneous experimental conditions are connected. CoRegNet is not vulnerable to the batch effect, can better implement integrated gene expression data, and can help to reveal non-linear gene relationships. To our knowledge, our study is the first to integrate more than 12,000 experiments from different labs with different goals to understand broad gene-gene associations. By integrating and analyzing all these experiments, we were able to generate a huge gene network revealing the most comprehensive relationship between genes at the RNA expression level. Our novel co-regulation network can be accessed via: http://CRN.liuzlab.org/

## METHOD

### Co-regulation Model

To integrate data from thousands of experiments, each with a small sample size and conducted under varied conditions, we first identify the genes that are changed due to a certain perturbation in individual experiment (Figure 1A). The perturbation is derived from genotype labels and phenotype changes given in the meta data. For samples where meta data label is not available, we used an unsupervised learning algorithm DASC to estimate a label[12]. The genes that are differentially expressed between two groups are called differentially expressed genes (DEGs); these DEGs are tabulated in a binary DEG matrix in which rows represent the genes and columns represent the experiments (Figure 1B).

**Figure 1.**
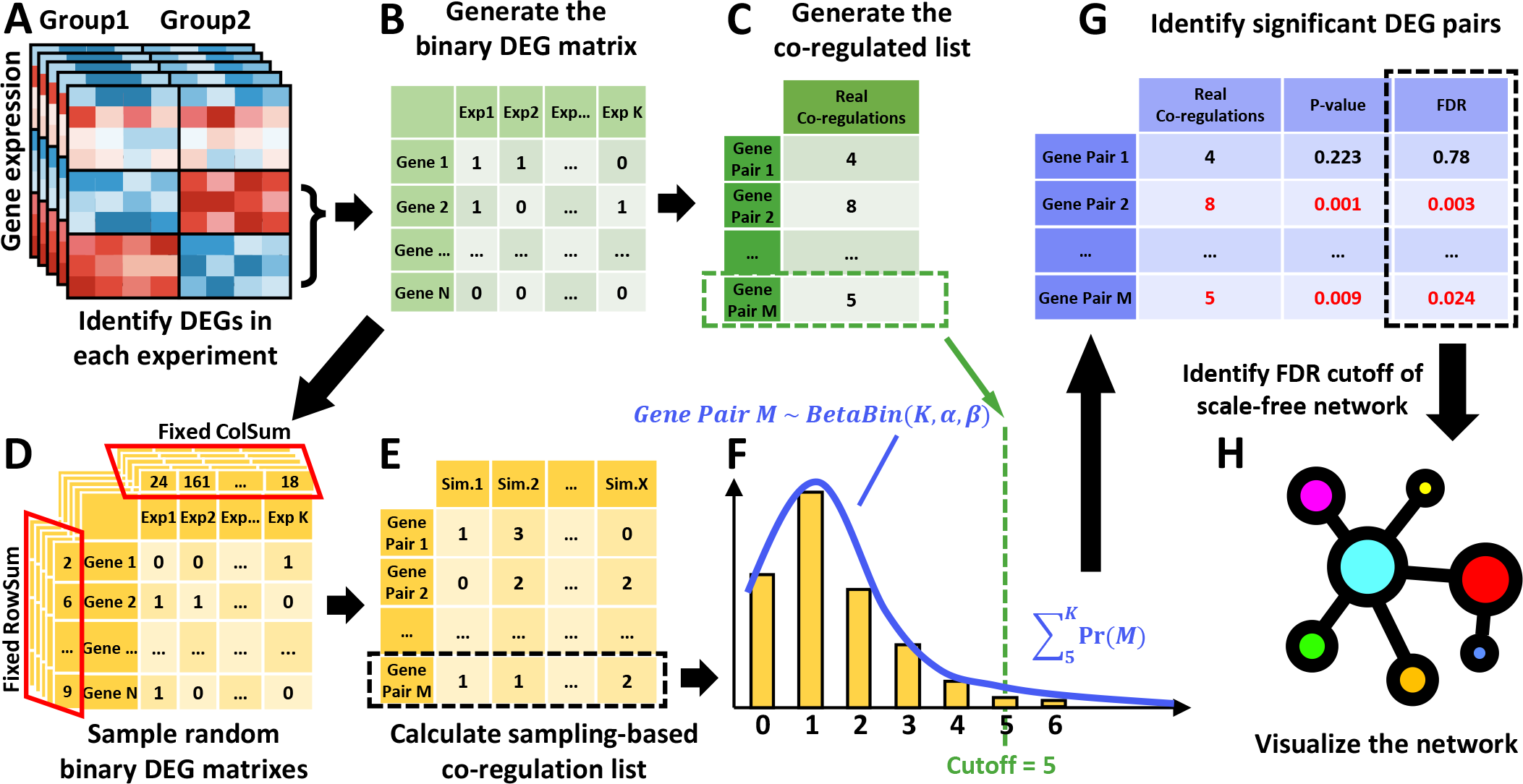
Co-regulation network (CoRegNet) framework. A) Identify co-regulated DEG pairs in each contrast. Each contrast contains two groups with a given label or a DASC label. B) Generate the binary DEG matrix with rows of genes and columns of contrasts. If the gene in the corresponding contrast is a DEG then marked as 1 otherwise 0. C) Generate the co-regulation list including all the gene pairs. The value in the list represents the number of contrasts each gene pair co-regulated. D) 1,000 random binary DEG matrices will be sampled with the same row and column sums as the true binary DEG matrix. E) 1,000 sampling-based co-regulation lists will be calculated based on each of the random binary DEG matrices. Each column represents a co-regulation list for each sampling, and each row is the random variable of a gene pair’s co-regulation F) Apply beta-binomial distribution to fit the random variable and take the true co-regulation times as the cutoff to identify significant DEG pairs. G) Calculate the p-value and FDR of each gene pair. H) Identify the FDR cutoff of a scale-free co-regulation network and visualize it.

Next, the number of experiments in which a gene pair is changed together are recorded in the co-regulation list (Figure 1C). The CoRegNet is designed to identify DEG pairs that are significantly changed together among different experiments. For example, we observed a *gene pair M* (Figure 1C) changed together *n*= 5 times out of a total of *K* experiments. To estimate the statistical significance of this observation, we need to calculate the distribution of the number of co-regulations over K experiment at random (a null model). We can treat the co-regulation in each experiment as a Bernoulli trial, where the probability of success (or co-regulation) *P*_*k*_ in each experiment is not fixed. Assuming *P*_*k*_ comes from a beta distribution, we can describe the random variable of *gene pair M* co-regulated in random cases as a beta-binomial distribution (Figure 1F)

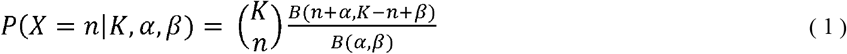

Where K is the total number of experiments, n is the number of co-regulated experiments, B is the Beta function, α and β are the parameters of Beta function.

The null distribution is generated by sampling random binary DEG matrices, each of which had the same sum of rows and sum of columns to accurately mimic the DEG probabilities (Figure 1D). In other words, our null hypothesis *H*_0_ is that (1) every single gene has the same chance of being a DEG as it does in the real data, and (2) each experiment provides the same number of DEGs as would actual data. Then, for each sampled binary DEG matrix, we calculated the sampling-based co-regulation list and combined every sampling together (Figure 1E). The row with gene pair M describes how many experiments it co-regulated in each random sampling, which should follow a beta-binomial distribution. We therefore used beta-binomial fitting on it and calculated its p-value 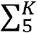 *Pr* (*M*) (Figure 1F). We used the same approach to calculate the p-value for all the gene pairs, and then calculated the FDR for them (Figure 1G). Based on the FDR, we chose the cutoff when the remaining significant DEG pairs can construct a scale-free gene network (Figure 1H).

### Evaluation of non-linearity between gene pairs

The CoRegNet revealed a novel set of gene pairs changed in the same direction in some conditions but changed in opposite directions in other conditions. In other words, a gene pair *X* and *Y* are concordant if and only if *X* goes up then *Y* goes up, or *X* goes down then *Y* goes down. A gene pair *X* and *Y* are discordant if and only if *X* goes up then Y goes down, or *X* goes down then *Y* goes up. In other words, if *X* and *Y* go in the same direction, we call this pair concordant. If *X* and *Y* go in opposite direction, we call this pair discordant. In order to evaluate the gene pair property that they are concordant in some conditions, and discordant in other conditions, we defined the non-linear gene pair score as:

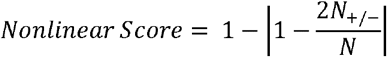

where N is the number of experiments in which a gene pair co-regulated and *N*_+/-_ corresponds to the number of experiments in which this gene pair discordantly co-regulated. The fundamental concept was to assign a non-linear score of 1 when the contrasts demonstrated equal proportions of concordant and discordant changes. Conversely, a score of 0 was assigned when all contrasts exhibited either entirely concordant or entirely discordant changes.

## RESULTS

### Non-linearly correlated gene pairs uncovered by CoRegNet

When applied our CoRegNet to Recount3 data, which contains more than 8,000 human studies, we found that a large number of gene pairs have non-linear correlation patterns due to heterogeneity. In Figure 2A, we showed a schematic diagram in which samples depicted in red and orange are concordantly correlated while samples depicted in blue and navy are discordantly correlated. Although we can discern their correlation separately, the overall correlation is very weak after combining all the samples, so pooling these data together would only ruin the co-expression pattern.

**Figure 2.**
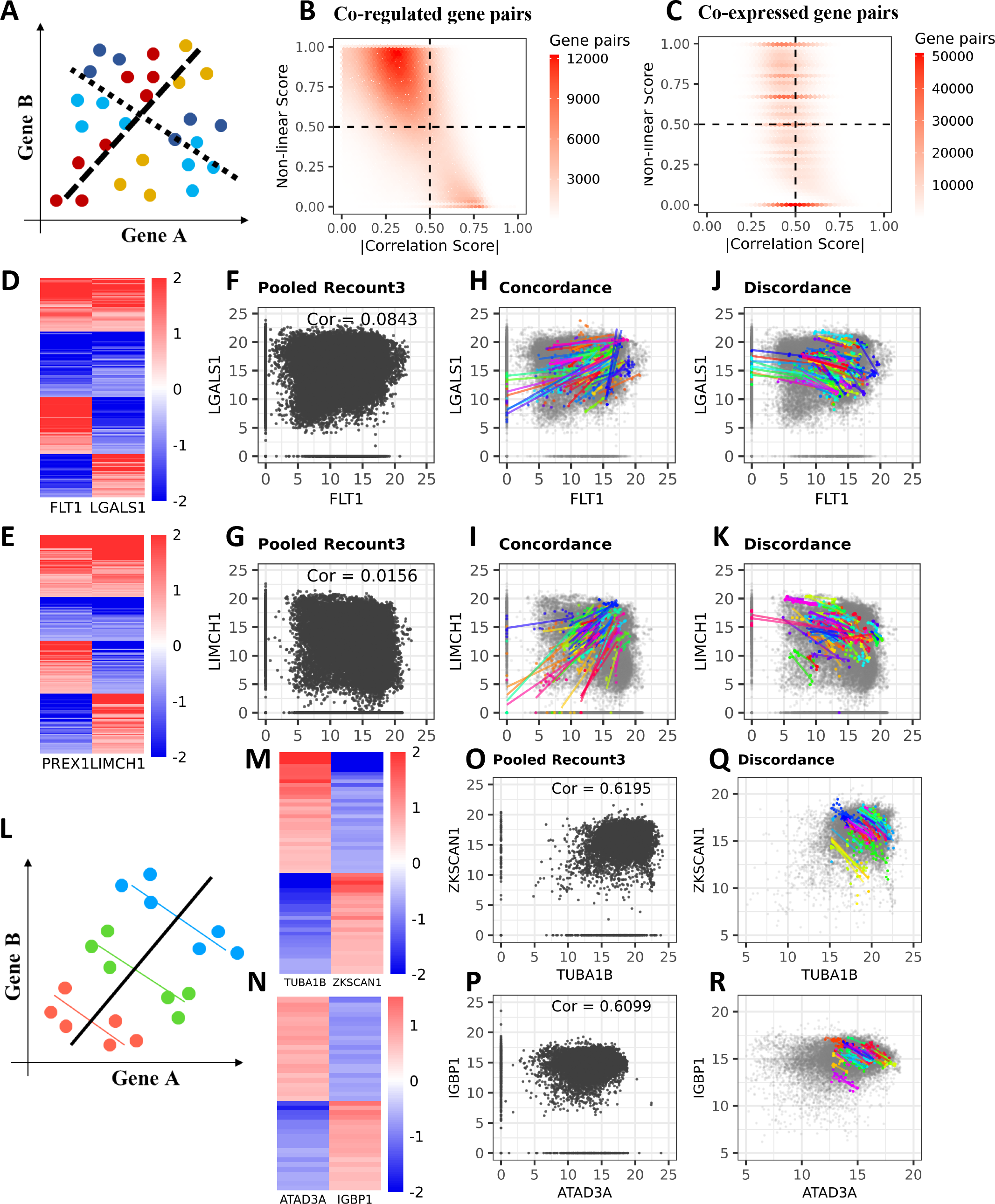
Co-regulation model can identify non-linear correlation and Simpson’s paradox in the integrated data. A) Schematic diagram of non-linear correlation. Samples in warm colors are concordantly correlated while samples in cold colors are discordantly correlated. B) Co-regulated gene pairs with low correlation scores tend to have higher non-linear scores in the integrated data. C) Co-expressed gene pairs tend to have a higher mean correlation score regardless of non-linear score. D,E) The log2FC heatmap of non-linear correlated gene pairs. F,G) The gene pair’s Pearson’s correlation score. H,I) Samples from contrasts that are concordantly co-regulated. J,K) Samples from contrasts that are discordantly co-regulated. L) Schematic diagram of Simpson’s paradox. M.N) The log2FC heatmap of negatively correlated gene pairs. O,P) The gene pair’s Pearson’s correlation score is positive. Q,R) Samples are only discordantly co-regulated in each contrast.

We calculated the non-linear score for the 3,000,000 significantly co-regulated gene pairs identified by our model out of the 360,000,000 total gene pairs. When plotted against the Pearson’s correlation, our results demonstrated that the gene pairs with high non-linear scores tend to have low Pearson’s correlations values (Figure 2B). In contrasts, gene pairs identified by WGCNA[13], tends to have high Pearson’s correlation and random non-linear scores, especially a large portion where non-linear score equals to 0 (Figure 2C). The reason why these co-regulated gene pairs have a low Pearson’s correlation is because they are non-linearly correlated among different contrasts (Figure 2B), which explains why a co-expression model like WGCNA cannot identify such gene-gene association.

We also showed two real examples in which the gene pairs were significantly co-regulated in many contrasts but were not always concordantly expressed together (Figure 2D-K). These two gene pairs are concordantly co-expressed in half of the contrasts and discordantly co-expressed in the other half (Figure 2D-E). However, in the pooled data, we can barely discern a linear correlation between two genes and the Pearson’s correlation score is close to 0 (Figure 2F-G). The Pearson’s correlation between FLT1 and LGALS1 is 0.0843, and the correlation between PREX1 and LIMCH1 is 0.0156. The co-expression model cannot identify either of the gene pairs although they are actually significantly co-regulated in 167 (FDR = 3.05 x 10^-12^) and 190 (FDR = 8.54 x 10^-2o^) contrasts, respectively. Both the concordant (Figure 2H-I) and discordant (Figure 2J-K) co-regulation are obvious when plotted separately.

Due to the context-dependent heterogeneity, gene pair correlations can be non-linear and hard to capture in the pooled data with traditional co-expression methodology. But our model successfully identified these gene-gene associations using co-regulated contrasts rather than co-expression patterns.

### Simpson’ s paradox in co-expression model

We also found that many of the gene-gene associations that WGCNA identified in pooled data were incorrect and that, in some of the gene-gene associations that did actually exist, the Simpson’s Paradox obfuscated their correlation trends[14].

Simpson’s paradox states that combining different groups of data together results in an overall correlation that is the opposite of each group’s true individual correlation[15]. When the CoRegNet showed that two genes were negatively correlated in every contrast, the co-expression model showed that the overall correlation was positive (Figure 2L). We used two examples to demonstrate that these two gene pairs are negatively correlated according to the log2FC heatmap (Figure 2M-N), but the overall correlation between TUBA1B and ZKSCAN1 and between ATAD3A and IGBP1 is ∼0.62. For both gene pairs, the Pearson’s correlation is positive, but if we look at the contrasts independently, they are both negatively correlated in all contrasts (Figure 2O-P).

Therefore, while the co-expression model may identify a significant connection between these genes, the genes ‘relationship can be completely reversed due to the Simpson’s paradox. One explanation for this is that the batch effect between different studies causes data bias so that, when integrating a large number of studies without batch correction, the correlation detected becomes the biased batch trend and not the gene signals. Overall, more than 40,000 gene pairs—90% of which are protein coding genes—are subject to the Simpson’s paradox.

### Biological function between linear and non-linear genes

To explore the biological differences between linear and non-linear genes, we selected the 5,861 most significant linear hub genes and the 3,849 most significant non-linear hub genes. We then compared these genes and the co-expression values of their neighboring genes in the GTEx dataset[16]. We found that the majority of linear genes identified in the Recount3 dataset also exhibited a positive correlation in the GTEx dataset, with a median correlation score of 0.5. However, surprisingly, the non-linearly correlated gene pairs had correlation scores ranging from -0.5 to 0.5, with most having very low correlation scores in the GTEx dataset (Figure 3A). These non-linear genes may often be overlooked by researchers due to their low correlation scores, but our findings indicate that these non-linear genes have distinct biological functions compared to linear genes. We created a Gene Ontology (GO) term tree, highlighting the terms enriched by both linear and non-linear genes. Our analysis revealed that linear genes significantly enrich 968 GO terms (FDR < 0.01), predominantly associated with *metabolic processes and cellular processes*. Conversely, non-linear genes enrich 902 GO terms (FDR < 0.01), with the most prominent terms relating to *developmental processes and biological regulation* (Figure 3B). We interpret that non-linear genes could exhibit positive or negative correlations with other genes under different conditions, an observation that aligns with the concept of biological regulation. Moreover, we found non-linear genes enrich terms associated with *system development*, particularly in the *generation of neurons*, suggesting that some non-linear genes may exert tissue-specific functions. In contrast, linear genes, which significantly enrich terms such as *DNA/RNA* metabolic process and cell cycle process, appear to perform more universal, non-tissue-specific functions. In addition to our GO analysis, we used the Human Phenotype Ontology[17] (HPO) to understand how linear and non-linear genes contribute to different diseases. Linear genes enriched 61 HPO terms (FDR < 0.05), and non-linear genes enriched 35 HPO terms (FDR < 0.05). Notably, we observed tissue-specific non-linear genes associated with nervous system diseases, as well as abnormalities of the head, neck, face, eye, and even patterns of inheritance. Conversely, linear genes were enriched in diseases associated with the immune system, blood, and growth (Figure 3C).

**Figure 3.**
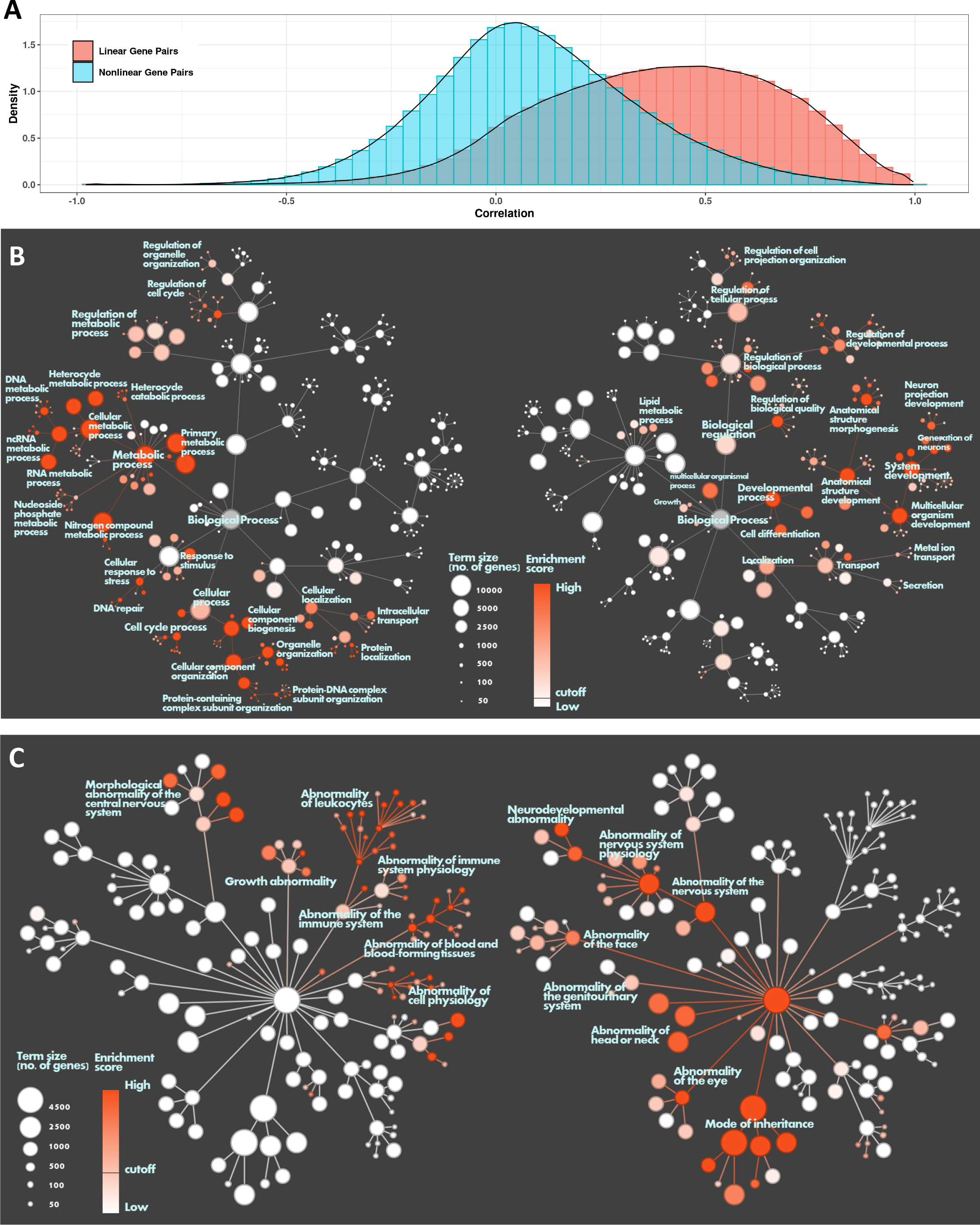
Comparison between linear and nonlinear genes: A) Histogram of Pearson’s correlation in GTEx data between top linear genes and non-linear genes. B, C) Left and right trees share the same topology of gene ontology(B) or human phenotype ontology(C), with nodes indicating terms and edges indicating hierarchical relations between terms. Node sizes represent the number of genes belong to a term. Node colors the enrichment significance, with cutoff where FDR < 0.01. The left tree is how linear genes are enriched, and the right tree is how non-linear genes are enriched. Nodes and modules with high enrichment significance are labeled.

### Co-regulation Network on Recount3 Data

For the purpose of analyzing and learning the biology behind the Recount3 co-regulation network, we used the recursive Louvain module detection method[18] on the Recount3 co-regulation network and identified 79 modules. Then, we generated a binary heatmap (Figure 4A) with rows of contrasts and columns of genes. The top color bar represents modules in the Recount3 co-regulation network while the left color bar indicates contrasts from different experiments. The black color indicates that the gene in the corresponding contrast is a DEG. We observed that genes in each module are co-regulated in a set of common contrasts, which come from various studies. This result means that genes co-regulated together are closely connected to each other in the network. It also begs the question whether, if a group of genes were co-regulated by the same set of contrasts, they share the same biological functions.

**Figure 4.**
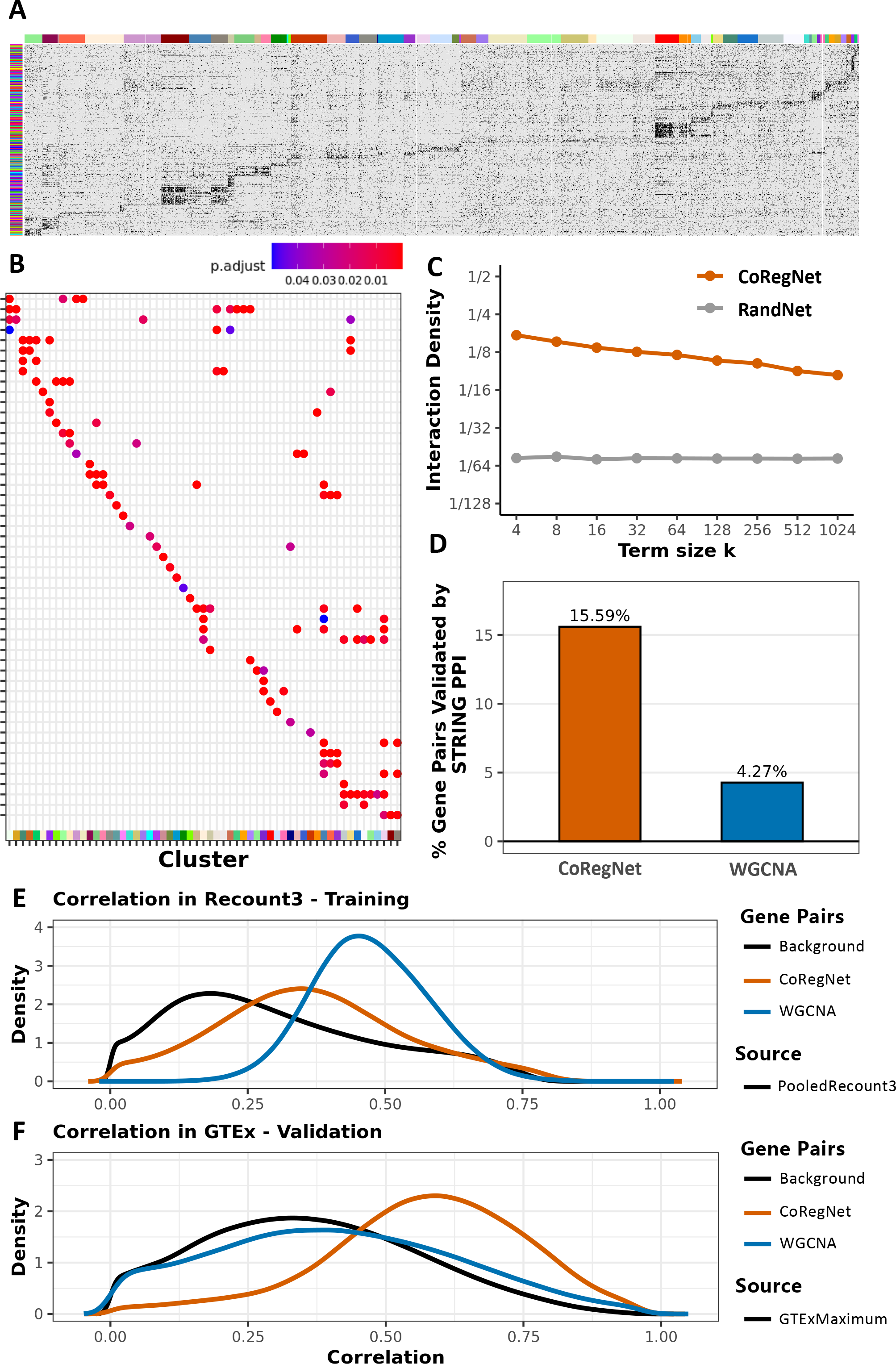
The co-regulation network on Recount3. A) Binary co-regulation heatmap with rows of contrasts and columns of genes. The left color bar represents different contrasts, and the top color bar represents genes in different modules. If a gene is a DEG in the contrast, it will be colored as black. B) GO enrichment comparison between different gene modules. C) GO term interaction density between Recount3 co-regulation network and random network. D) PPI validation between co-regulated gene pairs and co-expressed gene pairs. E) Comparison of Pearson’s correlation between co-regulated gene pairs and co-expressed gene pairs in Recount3 data. F) Validation by comparing Pearson’s correlation of co-expressed and co-regulated gene pairs from Recount3 in the independent GTEx data.

To answer this, we further investigated each module’s biological functions using the GO enrichment test and compared these modules’ common significant GO terms (Figure 4B). Multiple modules share some functions, like immune receptor activity and GTPase regulator activity, while other functions, like GDP binding and phosphatase activity, are unique to only one of the modules (Sup Figure 3A). In addition, we needed to confirm that these enriched GO terms derived from our co-regulation network are biologically meaningful and that a random network could not have the same quantity of enriched GO terms. We therefore compared our co-regulation network’s GO term interaction density with a random network that has the same genes and interaction density. We found that the co-regulation network has a considerably higher GO term interaction density than the random network (Figure 4C), which indicates that the gene-gene associations in the co-regulation network are biologically meaningful and not randomly assigned.

### Validation of Recount3 co-regulated genes

We assessed the performance of CoRegNet against the traditional co-expression method, WGCNA, using extensive datasets. We further validated our results by examining the protein-protein interactions between both co-regulated and co-expressed gene pairs, referencing the STRING database.[19]. We found that a higher percentage of gene pairs (Chi-Square p-value < 10^-20^) in the co-regulation network could be validated in the PPI database than could the co-expressed gene pairs (Figure 4D). PPI can detect about 15.6% of 3,000,000 co-regulated gene pairs and validate only 4.3% of 5,000,000 co-expressed gene pairs.

To ensure that the gene-gene associations we identified are generalizable—in other words, that the gene pairs found in Recount3 are also meaningful in another database—we used the gene pairs identified in Recount3 as training results and compared their correlations in a testing data set, which is GTEx data. We chose GTEx data because it provides a more homogenous background and ensures that all the samples are healthy human samples. We hypothesized that the gene-gene associations that Recount3 identified would be reproduced in the GTEx data. We found that in the Recount3 training data, the co-expressed gene pairs that WGCNA identified have a higher correlation score than co-regulated gene pairs (Figure 4 E). However, when we put them in GTEx testing data, we found that co-regulated gene pairs tend to have a higher correlation score and that co-expressed gene pairs only demonstrate a background correlation distribution in testing data (Figure 4F). These surprising results suggest that our CoRegNet does a better job finding the biologically meaningful gene pairs in the integrated data because a larger percentage of co-regulated gene pairs are validated by other independent databases.

### Co-regulation network on Recount2 Brain and MAD

In addition to the Recount3 co-regulation network, we also applied our CoRegNet to two smaller but domain-specific data sets: (1) the recount brain project[20] and (2) mouse Alzheimer’s disease (MAD). The recount brain project consists of 78 qualified studies with 93 total contrasts. These studies are all human brain-tissue-specific and contain ∼1,300 samples and 15,000 genes. The MAD dataset consists of 70 unique mouse studies and 138 contrasts and has ∼1,000 samples and 10,000 genes.

In Recount2 brain data, our CoRegNet identified 52,000 significantly co-regulated gene pairs (4,555 genes). The Louvain module detection method found 9 modules and for each module, we can clearly see that genes are co-regulated in a group of common contrasts (Figure 5A). Because these studies are all brain-region-specific, some gene modules shared many gene ontology (GO) terms in common (Figure 5B). These GO terms are related to nerves with diverse channel activities (Sup Figure 3B) such as ion channel activity. To further validate the biology behind the co-regulation network, we compared the Rcount2 brain co-regulation network with a random network with the same genes but different connections. The result showed that the co-regulation network has a higher GO term interaction density than the random network (Figure 5C). We also visualized the network in Figure 5D, in which the color of the genes corresponds to their module color.

**Figure 5.**
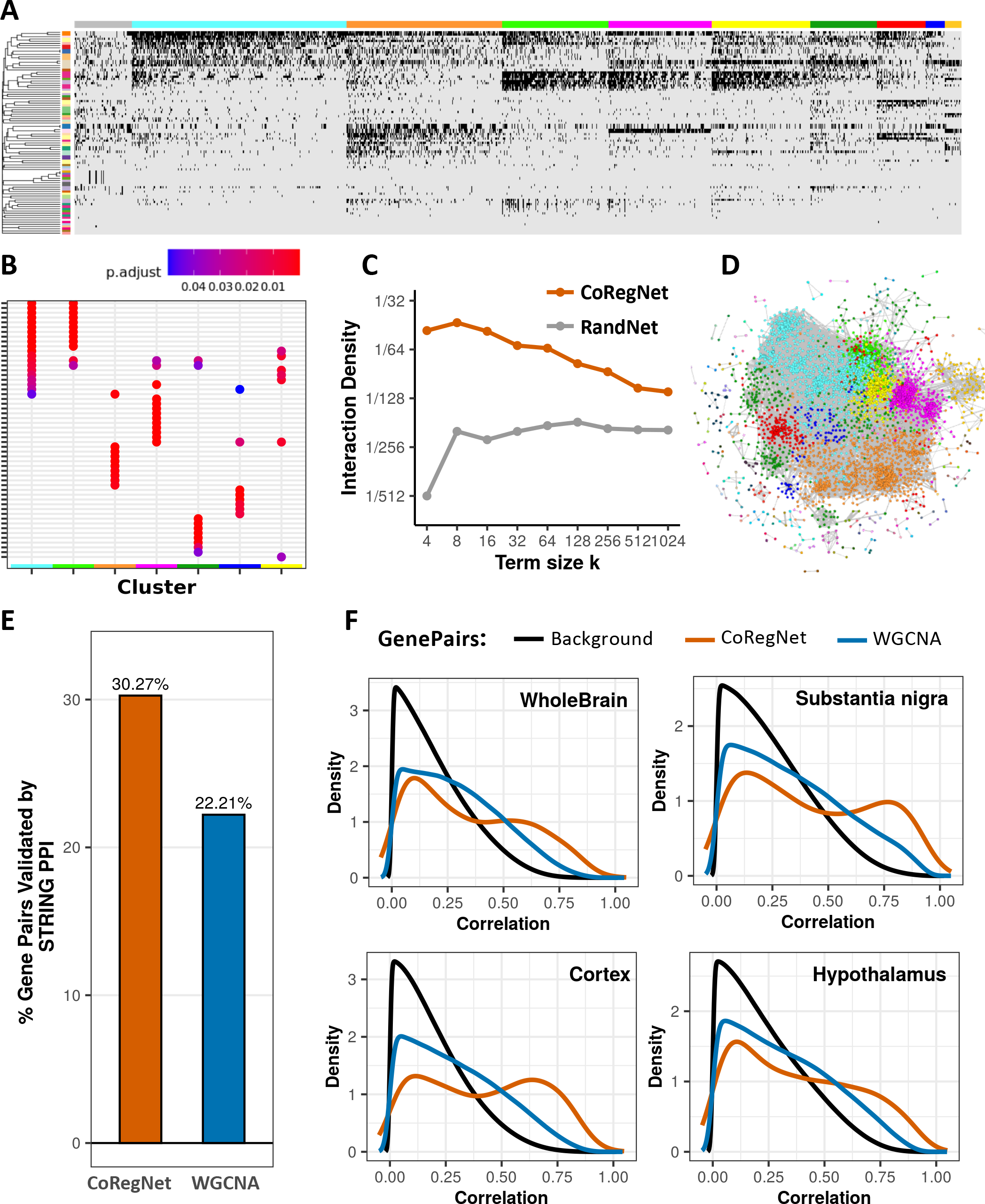
The co-regulation network on Recount2 Brain. A) Binary co-regulation heatmap with rows of contrasts and columns of genes. The left color bar represents different contrasts, and the top color bar represents genes in different modules. If a gene is a DEG in the contrast, it will be colored as black. B) GO enrichment comparison between different gene modules. C) GO term interaction density between Recount2 Brain co-regulation network and random network. D) Visualized Recount2 Brain co-regulation network. E) PPI validation between co-regulated gene pairs and co-expressed gene pairs. F) Validation by comparing Pearson’s correlation of co-expressed and co-regulated gene pairs from Recount2 Brain in the independent GTEx data.

Around 30% of the gene pairs recognized by CoRegNet were found to have protein-protein interactions according to the STRING database while only 22% of the gene pairs detected by WGCNA were validated by PPI data (Chi-Square p-value < 10^-20^) (Figure 5E). We also validated our gene pairs’ RNA seq data by GTEx brain data (Figure 5F). Regarding background distribution, most gene pairs in the brain GTEx had a low correlation score, while gene pairs identified by WGCNA in Recount2 Brain had similar background distribution as the brain GTEx. For CoRegNet, however, a bimodal distribution showed that, though many gene pairs still had a low correlation score, half of the co-regulated gene pairs have a high correlation score in GTEx brain data. Gene pairs identified by the CoRegNet in Recount2 Brain are generally better validated than those identified by WGCNA in both the GTEx data and STRING database.

In the second small case study, we used the CoRegNet on the MAD dataset to successfully identify 201,454 significantly co-regulated gene pairs (7,199 genes). The Louvain module detection approach found 10 modules in the network, and genes co-regulated in common contrasts were clustered together (Sup Figure 4A). Because the MAD studies are more domain-specific—they are all related to neurodegenerative diseases in mice—the GO term enrichment results showed that some modules share many common GO terms (Sup Figure 4B). These enriched GO terms were also related to channel activities and channel bindings (Sup Figure 3C). We performed the GO term interaction density test, which demonstrated that the MAD co-regulation network still has a higher interaction density than the random network. This in turn suggests that there is agreement between the co-regulation network and GO terms (Sup Figure 4C). Finally, we visualized the MAD co-regulation network, in which the color of genes corresponds to their module color (Sup Figure 4D). The orange and turquoise modules shared many common GO terms, and they are closely connected to each other in the network.

Based on our PPI validation, genes identified as co-expressed via WGCNA exhibited a higher PPI rate compared to those identified as co-regulated. Although the difference was statistically significant (Chi-Square p-value < 10^-20^), the validation percentages between the two groups were closely matched, with a disparity of less than 1% (Sup Figure 4E). Also, we used GTEx data to perform RNA validation and borrowed disease oriented ROSMAP data[21]. We found that co-expressed genes from WGCNA could not be accurately validated in either the GTEx or ROSMAP data because the correlation distribution still follows the background distribution in both independent datasets. The co-regulated genes ‘correlation distribution in MAD, however, had a higher average correlation score (Sup Figure 4F), indicating that co-regulated gene pairs are still better validated than co-expressed gene pairs.

### Example of gene function prediction using the Recount3 co-regulation network

Interferon-induced protein with tetratricopeptide repeats (IFIT) genes are the well-known interferon-stimulated genes (ISG)[22]. This family contains four genes: IFIT1, IFIT2, IFIT3, and IFIT5. In our Recount3 co-regulation network, we successfully identified all four IFIT genes and ISG15 in the same sub-network (Figure 6A). We also found that several other genes in this subnetwork shared similar disease-related functions. According to the OMIM (Online Mendelian Inheritance in Man) database, ISG15, IRF9, and STAT1 are related to immunodeficiency. These three genes belong to the ISG15 conjugation pathway, in which IRF9 interacts with phosphorylated STAT1 and STAT2 to form the ISGF3 complex; it then recognizes the interferon-sensitive response elements that promote ISG15 and its conjugation enzymes to induce ISG15 expression[23]. This same study confirmed that ISG15 plays an important role in the host antiviral response[23]. We also found that the IFIH1 and DDX58 genes are associated with Singleton-Merten syndrome, a multi-system innate immune disorder[24,25]. Reactome shows that they play a key role in DDX58/IFIH1-mediated induction of the interferon-alpha/beta pathway, which is also part of the innate immune system[26]. Further, SP110 mutations associate with hepatic venoocclusive disease with immunodeficiency, a rare form of severe combined immune deficiency[27]. Genes SAMD9 and SAMD9L in this sub-network are both related to monosomy 7 myelodysplasia and leukemia syndrome 1. Though these two genes are not immunodeficiency related, they are antiviral factors that play crucial roles in the innate immune system’s defense against poxviruses[28].

**Figure 6.**
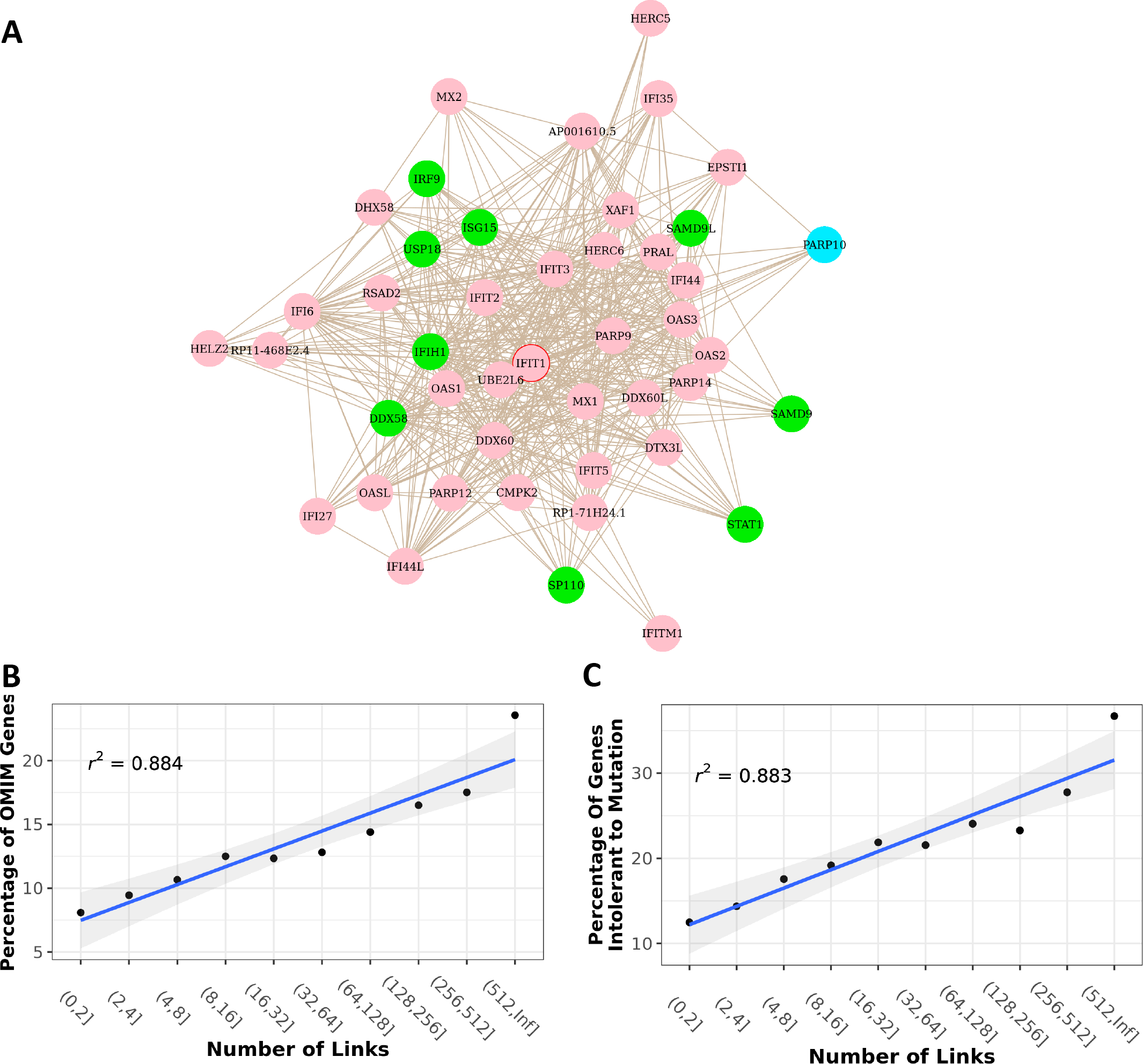
A) The IFIT family subnetwork. The green genes have a known OMIM phenotype. The blue genes lack known OMIM phenotypes but are recorded as pathogenic in ClinVar. The pink genes lack known OMIM phenotypes and are recorded as non-pathogenic in ClinVar. B) The proportion of OMIM gene neighbors is positively correlated with the total number of neighbors a gene has. C) The proportion of lethal gene neighbors is positively correlated with the total number of neighbors a gene has.

Based on the findings detailed above, we are optimistic that we will be able to predict known genes ‘unknown gene functions. In the IFIT family subnetwork, ClinVar records the PARP10 gene as a pathogenic gene not yet linked to any specific disease phenotype[29]. But, because many genes in this subnetwork are immunodeficiency-related, we believe that PARP10 has a higher likelihood of being associated with immunodeficiency. As we expected, multiple studies have confirmed PARP10’s critical role in regulating innate immune responses[30] and host-pathogen interactions[31].

### Increased essentiality of hub genes in the Recount3 Co-regulation network

The Recount3 co-regulation network consists of significantly co-regulated gene pairs. Interestingly, we found that when we increase the FDR threshold, the network becomes increasingly scale-free (Sup Figure 5A), which means that a small number of genes connect with many neighbors while a large number of genes connect with very few neighbors. We analyzed these hub genes in the scale-free co-regulation network and found that they are more likely to be disease- and lethality-related.

We ranked each gene based on its number of links and correlated this with the disease label from the OMIM database. We found that the likelihood of a gene being disease-related is positively correlated with the gene’s node degree in the network (the number of neighbors a gene has) (Figure 6B). Such disease gene frequency is stronger than the one reported by Barabási et al[32]. According to the scale-free co-regulation network, more than half the genes contain fewer than 8 links and 13% of them are disease-related genes. And while only 2% of the genes have over 512 links, ∼30% of them are disease related. We borrowed the mutation information from GnomAD (Genome Aggregation Database) and analyzed the relationship between the genes ‘degrees and mutation intolerance (Figure 6C). The result showed that the hub genes are less likely to tolerate loss-of-function (LoF) mutations. Around 80% of genes with fewer than 4 links tolerate LoF alleles. In other words, losing one gene copy is likely not lethal. In contrast, ∼40% of genes with more than 512 links cannot tolerate LoF alleles. Besides, we noticed that these disease-related genes consist of both protein coding and non-protein coding genes. We found a strong correlation between the proportion of protein coding genes and the gene’s node degree (Sup Figure 6A), however, when we only focus on the correlations between protein coding disease-related genes and the node degree, the correlations are obvious yet less significant (Sup Figure 6B-C). Therefore, we believe that the central genes in the Recount3 co-regulation network are more functional-essential genes. They have a stronger connection with disease and a greater impact on vitality.

## CONCLUSION

Our study provides a new method to reveal gene-gene associations by integrating a large amount of bulk RNA sequencing data without doing batch correction. Besides, our model is able to identify novel gene pairs that are non-linearly correlated between different experiments. We investigated our co-regulation network’s property and found that the hub genes are more likely to be essential and disease-related genes. Thus, we believe that our Recount3 co-regulation demonstrates a more comprehensive gene-gene relationship which can better assist biologists in discovering unknown gene functions and potential disease relations.

Given the many studies that collect bulk RNA sequencing data, we are the first to utilize as many studies as we can to generate the gene network. Meanwhile, our model is not limited to 12,000 experiments because our CoRegNet is not influenced by batch effect, which is the main challenge when integrating data from numerous experiments. The batch effect is a difficult obstacle to overcome using a reliable statistical algorithm. We compared the histogram of Pearson’s correlation scores of all gene pairs between batch-corrected and non-batch-corrected GTEx data (Sup Figure 7A-B) and found that, for some tissues, including brain, kidney, and blood, the peak is completely biased before batch correction. Although the distributions in other tissues have their peaks in low correlation regime, the density is still different, and there are more gene pairs with correlations close to zero after batch correlation. Thus, batch effect is significant in pooled GTEx data and can influence correlation distribution. We analyzed the correlation distribution and observed a biased peak in all three of the pooled data we used (Sup Figure 7C); these results are similar to the GTEx data before batch correction (Sup Figure 7A). We therefore believe that the batch effect is likely to occur when integrating expression data from numerous studies.

After we thoroughly analyzed the global gene-gene correlations of significantly co-regulated gene pairs, we highlighted two major findings. First, the correlation of the same gene pair can be reversed when treated with different experimental conditions. Due to such heterogeneity, when integrating various large-scale studies, a combination of concordant correlation and discordant correlation will lead to a disordered and non-linear relationship. Such non-linear correlation usually has a low Pearson’s correlation score which co-expression models will struggle to identify. Our study verified that the gene pairs with low correlation scores in the integrated data that our CoRegNet identifies will have higher correlation scores in the independent GTEx data; this means that those gene-gene correlations are biologically meaningful and deserve scientific attention because of their context-dependent relationship. We also noticed that even though some gene pairs have high positive Pearson’s correlation scores in the integrated data, the true correlations between those gene pairs are negative. This phenomenon is called Simpson’s paradox and we believe it may be caused by batch effect between experiments. Thus, the correlation model captures the global batch trend while the true local correlations of every experiment are actually negative. Our CoRegNet can successfully recognize non-linear correlation trends and is not confused by Simpson’s paradox.

We also examined our comprehensive Recount3 co-regulation network and found that linear genes and non-linear genes perform fundamentally distinct roles across various biological processes and diseases. Meanwhile, in the scale-free network, the hub genes have a higher proportion of disease-related genes and function-essential genes. This met our expectations because we believe that those important genes are more likely to interact with different genes under different conditions. To our surprise, our results exceed the classical PPI hub gene results demented by the Barabási et al[32]. We also used the several subnetworks to show that functionally related genes are connected to each other.

Apart from the huge amount of Recount3 data, we also demonstrated the power of CoRegNet on two small but domain-specific datasets (Recount2 Brain and MAD). In these applications, our CoRegNet not only recognized highly correlated gene pairs in the pooled data, but also genes that have low correlation scores because they are non-linearly correlated in the integrated data. We showed that, in all three cases, other independent databases better validated our co-regulated gene pairs than co-expressed gene pairs. We also investigated the biology behind our co-regulation network. Compared with the modules identified in the Recount3 network, the GO enrichment test suggests that modules in the Recount2 Brain and in MAD share more common GO terms. As Recount2 Brain and MAD are domain-specific studies, those genes are all detected in human brain-region-based or mouse-neurodegenerative-disease studies. So, it is unsurprising that the different modules share many similar functions in these two small datasets.

With the current model, treating millions of gene pairs is prohibitively time-consuming. Although generating a sampling-based binary DEG matrix is fast, fitting beta-binomial distribution to every single gene pair is not. Because calculating the random variable of a gene pair that will be co-regulated is an NP problem, we used a sampling and fitting approach in the model. Now that the Recount3 co-regulation network is available to the community, we hope and expect that it will drive new scientific insights, reveal novel gene connections, and predict unknown gene functions.

### Key Points

- Integration of extensive experiments performed under diverse conditions presents a formidable challenge. In this study, we introduce a statistical model tailored to discern significantly co-regulated gene pairs by amalgamating data from thousands of experiments. This method offers a more comprehensive gene-gene association network. Within the Recount3 co-regulation network, we observed a noteworthy trend: hub genes are predominantly associated with diseases and exhibit lethal characteristics.
- CoRegNet obviates the need for batch corrections even when assimilating a vast array of experiments. Our findings indicate that CoRegNet surpasses the performance of the widely used WGCNA in identifying gene-gene associations. Importantly, CoRegNet remains uninfluenced by the Simpson’s paradox. While conventional co-expression models might identify a gene pair as being overall positively correlated, CoRegNet can astutely highlight instances where, within individual sub-experiments, these genes exhibit negative correlations.
- CoRegNet can reveal non-linearly correlated gene pairs. The dynamics of these non-linear gene pair relationships are intriguing, as their correlation patterns are mutable and can shift under varying conditions. Delving deeper, we discovered distinct functional disparities between linear and non-linear genes. While linear genes predominantly participate in functions like metabolic processes, the non-linear genes show a stronger affiliation with biological regulation and developmental processes.

## Supporting information

Sup Animation1

## CODE AND DATA AVAILABILITY

Our co-regulation model R package is available from the GitHub repository: https://github.com/Jiasheng-Wang/CoRegNet

The datasets analyzed during the current study are available:

-https://rna.recount.bio/, Recount3

-http://research.libd.org/recount-brain/, Recount2 Brain

-https://doi.org/10.7303/syn2580853, MAD

## ACKNOWLEDGMENTS

We thank Alexander J. Trostle, Hyun-hwan Jeong, Chaozhong Liu for their contributions; Sasidhar Pasupuleti for his help managing the network website.

## Author Biographies

**Jiasheng Wang** is a PhD candidate at Jan and Dan Duncan Neurological Research Institute at Texas Children’s Hospital, Houston, Graduate Program in Quantitative and Computational Biosciences in Baylor College of Medicine and Department of Pediatrics in Baylor College of Medicine. His research interests are focused on Bioinformatics, Statistical Modeling, Data Mining, Machine Learning and Deep Learning.

**Ying-Wooi Wan** is an assistant professor at Jan and Dan Duncan Neurological Research Institute at Texas Children’s Hospital and Department of Molecular and Human Genetics in Baylor College of Medicine. Her research interests are focused on Bioinformatics, Statistical Modeling and Machine Learning.

**Rami Al-Ouran** is an assistant professor at Al Hussein Technical University. His research interests are focused on Bioinformatics, Machine Learning, Data Mining, and Computational Regulatory Genomics

**Meichen Huang** is a graduate student at Jan and Dan Duncan Neurological Research Institute at Texas Children’s Hospital and Department of Neurology in Baylor College of Medicine. Her research interests are focused on Bioinformatics, Statistical Modeling and Neurodegenerative Diseases.

**Zhandong Liu** is a professor at Jan and Dan Duncan Neurological Research Institute at Texas Children’s Hospital, Graduate Program in Quantitative and Computational Biosciences in Baylor College of Medicine and Department of Pediatrics in Baylor College of Medicine. His research interests are focused on Bioinformatics, Statistical Modeling, Graphical Modeling, Machine Learning, Deep Learning, Data mining, Genomics, Multi-omics and Neurodegenerative Diseases.

## Author notes

*J*.*W. and Z*.*L. led the research. J*.*W developed the model and pipeline. J*.*W*., *Y*.*-W*.*W. and R*.*A*.*-O. analyzed the networks. J*.*W*., *Y*.*-W*.*W. and Z*.*L. wrote the paper. M*.*H. designed the UI of the website*.

## Reference

1. Barrett T, Wilhite SE, Ledoux P, et al. NCBI GEO: archive for functional genomics data sets—update. Nucleic Acids Res 2013; 41:D991–D995

2. Kolesnikov N, Hastings E, Keays M, et al. ArrayExpress update—simplifying data submissions. Nucleic Acids Res 2015; 43:D1113–D1116

3. Ritchie ME, Phipson B, Wu D, et al. limma powers differential expression analyses for RNA-sequencing and microarray studies. Nucleic Acids Res 2015; 43:e47–e47

4. Zhang Y, Parmigiani G, Johnson WE. ComBat-seq: batch effect adjustment for RNA-seq count data. NAR Genom Bioinform 2020; 2:qaa078

5. Furlotte NA, Kang HM, Ye C, et al. Mixed-model coexpression: calculating gene coexpression while accounting for expression heterogeneity. Bioinformatics 2011; 27:i288–i294

6. Song L, Langfelder P, Horvath S. Comparison of co-expression measures: mutual information, correlation, and model based indices. BMC Bioinformatics 2012; 13:328

7. Jie T H HJ A HD, et al. ADAGE-Based Integration of Publicly Available Pseudomonas aeruginosa Gene Expression Data with Denoising Autoencoders Illuminates Microbe-Host Interactions. mSystems 2016; 1:10.1128/msystems.00025-15

8. Tan J, Doing G, Lewis KA, et al. Unsupervised Extraction of Stable Expression Signatures from Public Compendia with an Ensemble of Neural Networks. Cell Syst 2017; 5:63–71.e6

9. Zhou W, Altman RB. Data-driven human transcriptomic modules determined by independent component analysis. BMC Bioinformatics 2018; 19:327

10. Taroni JN, Grayson PC, Hu Q, et al. MultiPLIER: A Transfer Learning Framework for Transcriptomics Reveals Systemic Features of Rare Disease. Cell Syst 2019; 8:380–394.e4

11. Bonett DG, Wright TA. Sample size requirements for estimating pearson, kendall and spearman correlations. Psychometrika 2000; 65:23–28

12. Yi H, Raman AT, Zhang H, et al. Detecting hidden batch factors through data-adaptive adjustment for biological effects. Bioinformatics 2018; 34:1141–1147

13. Langfelder P, Horvath S. WGCNA: an R package for weighted correlation network analysis. BMC Bioinformatics 2008; 9:559

14. Blyth CR. On Simpson’s Paradox and the Sure-Thing Principle. J Am Stat Assoc 1972; 67:364–366

15. Wang B WU P, Kwan B, et al. Simpson’s Paradox: Examples. Shanghai Arch Psychiatry 2018; 30:139–143

16. Lonsdale J, Thomas J, Salvatore M, et al. The Genotype-Tissue Expression (GTEx) project. Nat Genet 2013; 45:580–585

17. Köhler S, Gargano M, Matentzoglu N, et al. The Human Phenotype Ontology in 2021. Nucleic Acids Res 2021; 49:D1207–D1217

18. Blondel V, Guillaume J-L, Lambiotte R, et al. Fast Unfolding of Communities in Large Networks. Journal of Statistical Mechanics Theory and Experiment 2008; 2008:

19. Szklarczyk D, Gable AL, Lyon D, et al. STRING v11: protein–protein association networks with increased coverage, supporting functional discovery in genome-wide experimental datasets. Nucleic Acids Res 2019; 47:D607–D613

20. Razmara A, Ellis SE, Sokolowski DJ, et al. recount-brain a curated repository of human brain RNA-seq datasets metadata. BioRxiv 2019; 618025

21. Bennett DA, Buchman AS, Boyle PA, et al. Religious Orders Study and Rush Memory and Aging Project. Journal of Alzheimer’s Disease 2018; 64:S161–S189

22. Pidugu VK, Pidugu HB, Wu M-M, et al. Emerging Functions of Human IFIT Proteins in Cancer. Front Mol Biosci 2019; 6:

23. Perng Y-C, Lenschow DJ. ISG15 in antiviral immunity and beyond. Nat Rev Microbiol 2018; 16:423–439

24. Ferreira CR, Crow YJ, Gahl WA, et al. DDX58 and Classic Singleton-Merten Syndrome. J Clin Immunol 2019; 39:75–80

25. Jang M-A, Kim EK, Now H, et al. Mutations in <em>DDX58</em>, which Encodes RIG-I, Cause Atypical Singleton-Merten Syndrome. The American Journal of Human Genetics 2015; 96:266–274

26. Jassal B, Matthews L, Viteri G, et al. The reactome pathway knowledgebase. Nucleic Acids Res 2020; 48:D498–D503

27. Wang T, Ong P, Roscioli T, et al. Hepatic veno-occlusive disease with immunodeficiency (VODI): First reported case in the U.S. and identification of a unique mutation in Sp110. Clinical Immunology 2012; 145:102–107

28. Peng S, Meng X, Zhang F, et al. Structure and function of an effector domain in antiviral factors and tumor suppressors SAMD9 and SAMD9L. Proceedings of the National Academy of Sciences 2022; 119:e2116550119

29. Landrum MJ, Chitipiralla S, Brown GR, et al. ClinVar: improvements to accessing data. Nucleic Acids Res 2020; 48:D835–D844

30. Zhu H, Tang Y-D, Zhan G, et al. Corrigendum: The critical role of PARPs in regulating innate immune responses. Front Immunol 2023; 14:

31. Fehr AR, Singh SA, Kerr CM, et al. The impact of PARPs and ADP-ribosylation on inflammation and host–pathogen interactions. Genes Dev 2020; 34:341–359

32. Jeong H, Mason SP, Barabási A-L, et al. Lethality and centrality in protein networks. Nature 2001; 411:41–42

