## Supplemental Figures for "CoRegNet: Unraveling Gene Co-regulation Networks from Public RNA-Seq Repositories Using a Beta-Binomial Statistical Model"

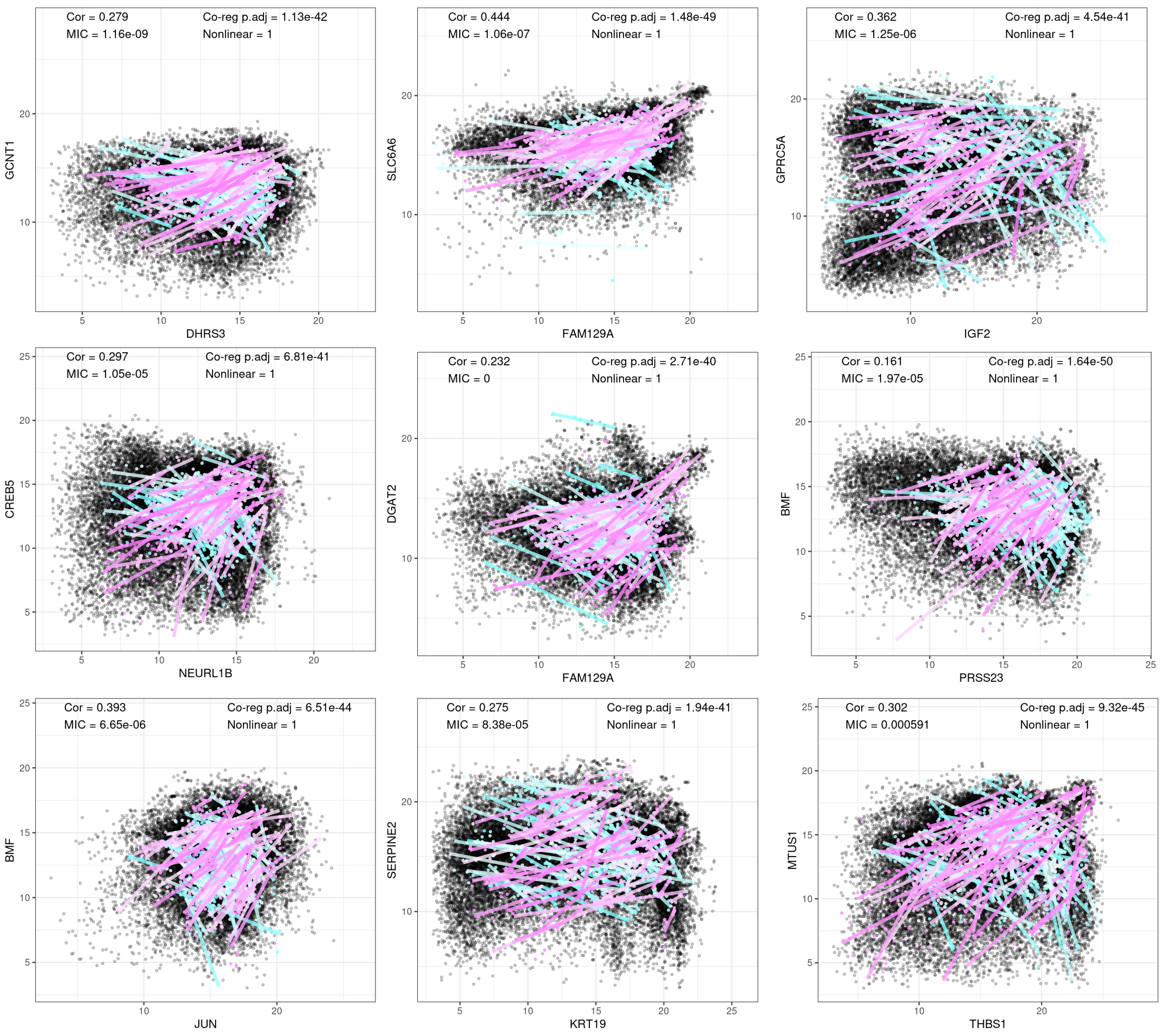


Supplementary Figure 1: Nine examples that non-linearly correlated gene pairs can be robustly identified by co-regulation model, but their mutual information coefficient (MIC) are small. In each sub-plot, the upper left corner shows the Pearson's correlation score and MIC of the gene pair, and the upper right corner shows the co-regulation significance (FDR) and the non-linear score of the gene pair. The black points represent all samples from Recount3. Each red line represents a contrast that the gene pair is concordantly changed. Each blue line represents a contrast that the gene pair is discordantly changed.

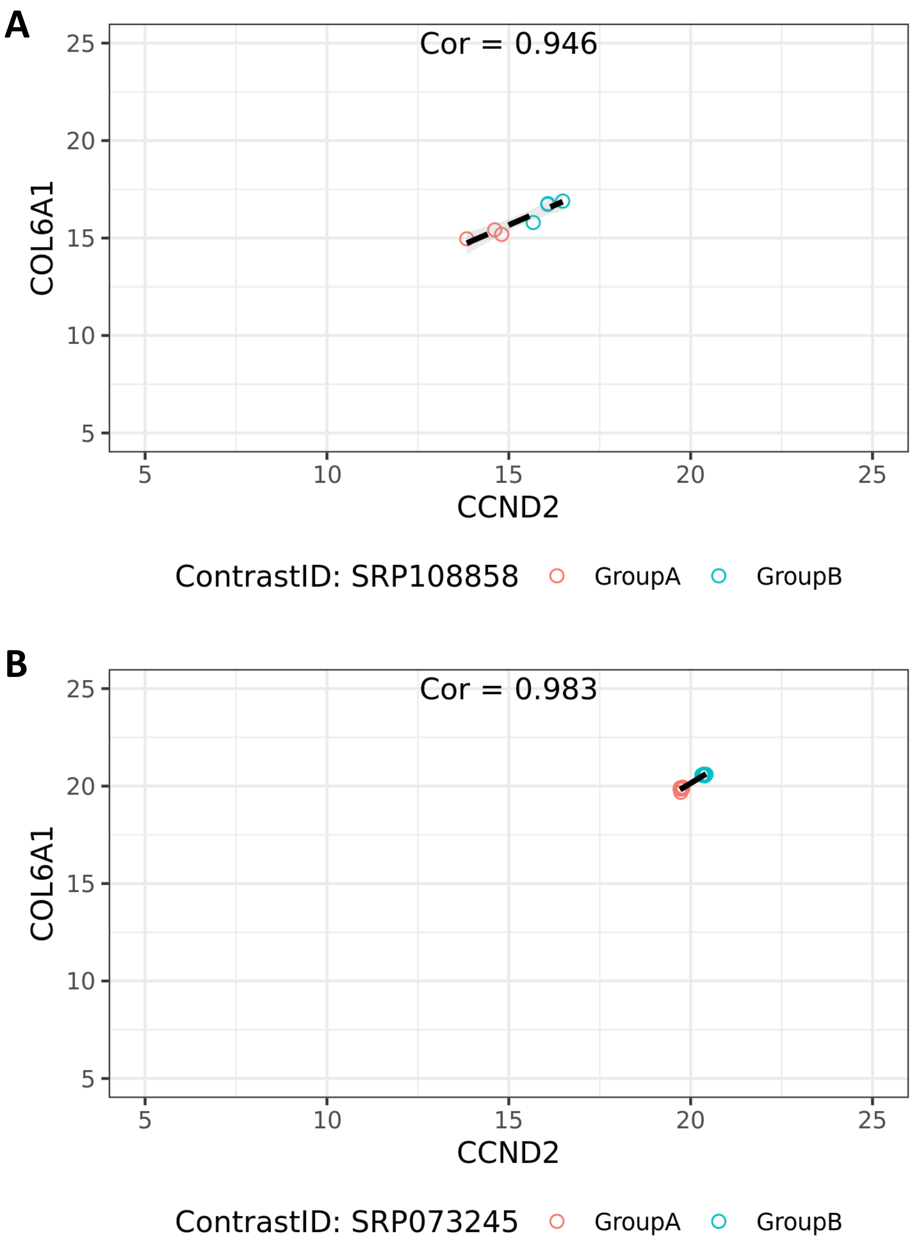


Supplementary Figure 2: The data quality influences the Pearson's correlation score. A) The expected situation that samples are not overlapped with each other. B) 6 samples in group A and 6 samples in group B have very similar expression so that they overlap with each other. Even the correlation score is high, such correlation is not based on 12 different samples. It's more like a linear regression on only two points.

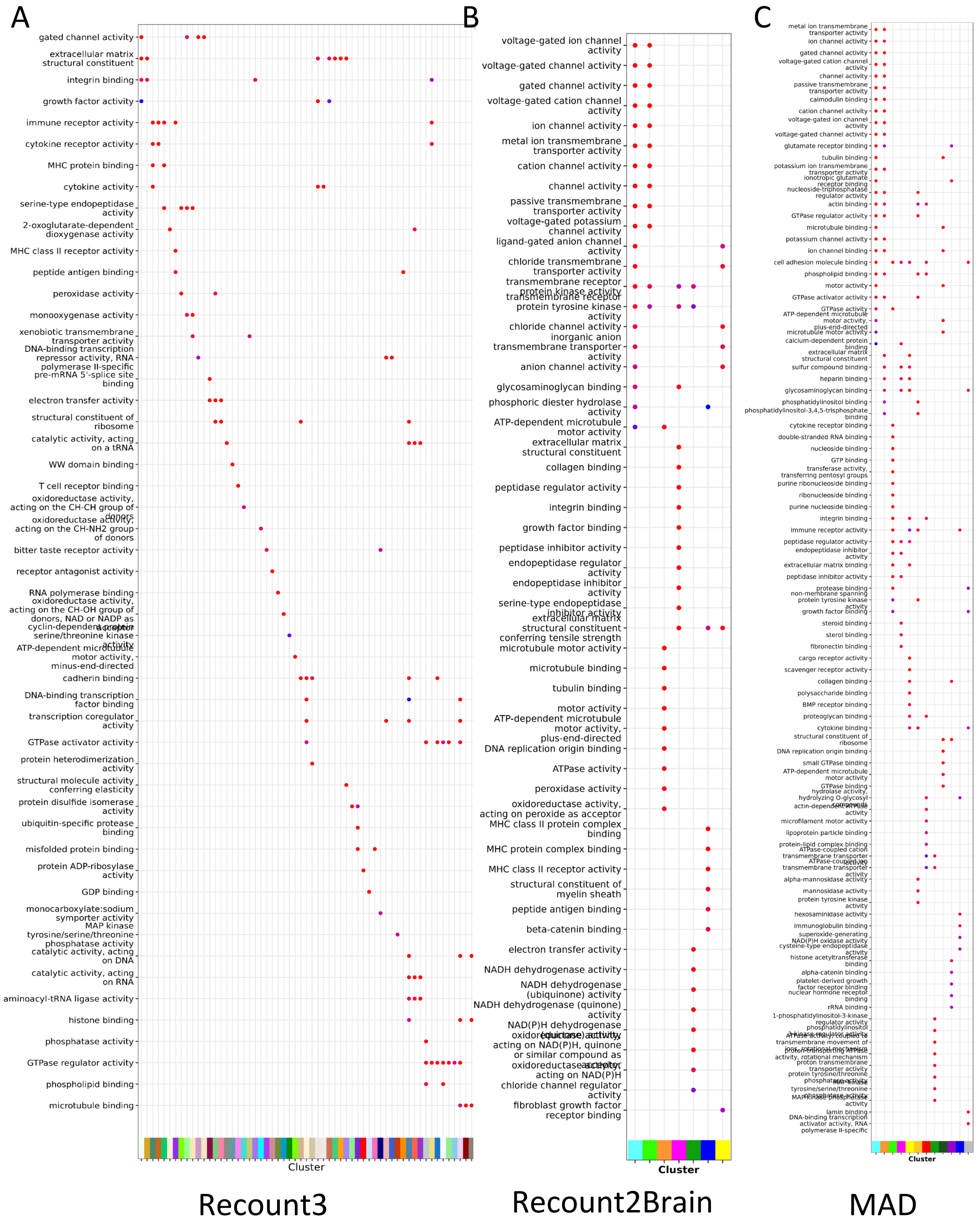


Supplementary Figure 3 Detailed GO enrichment comparison between different gene modules.

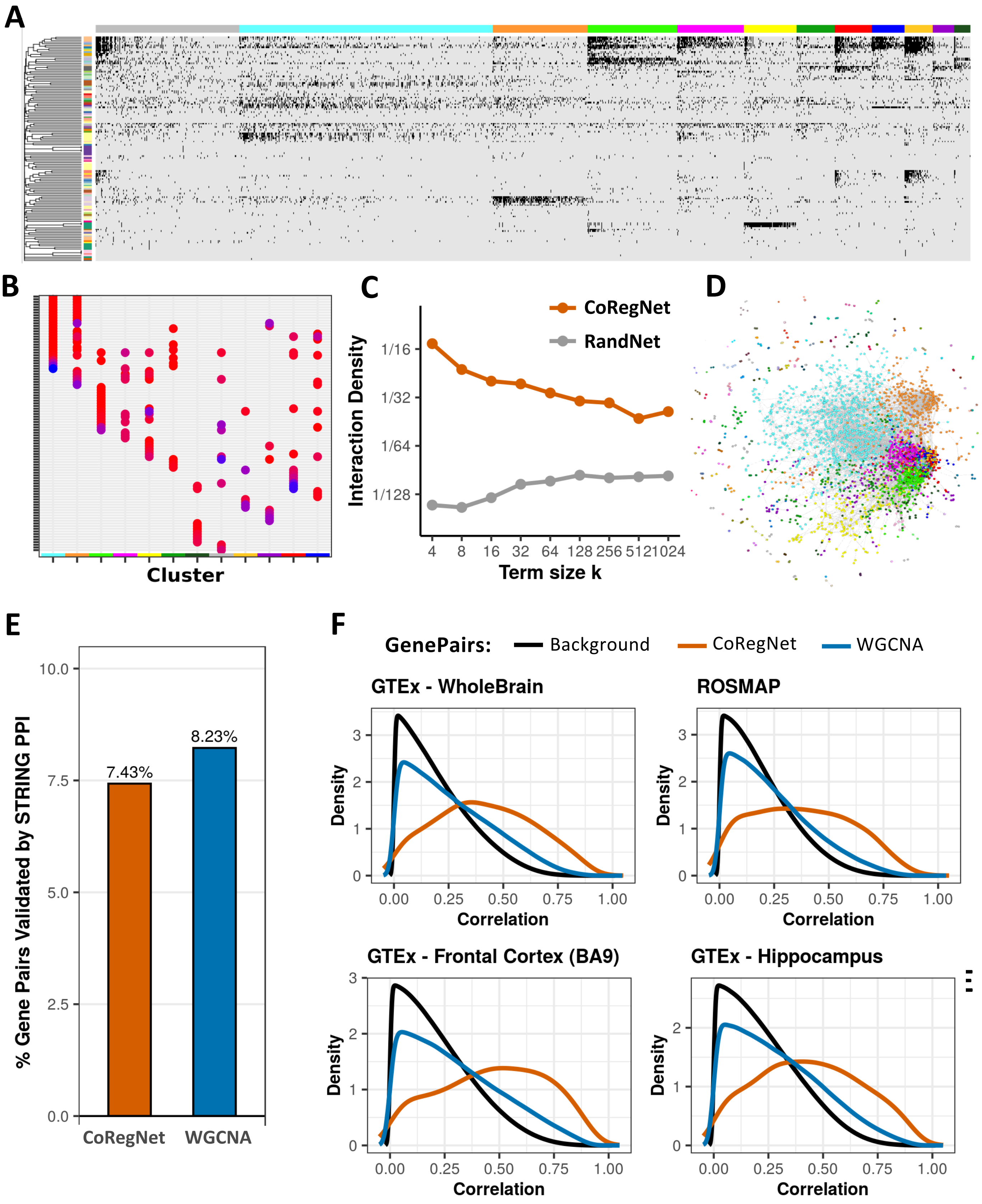


Supplementary Figure 4. The co-regulation network on MAD. A) Binary co-regulation heatmap with rows of contrasts and columns of genes. The left color bar represents different contrasts, and the top color bar represents genes in different modules. If a gene is a DEG in the contrast, it will be colored as black. B) GO enrichment comparison between different gene modules. C) GO term interaction density between MAD co-regulation network and random network. D) Visualized MAD co-regulation network. E) PPI validation between co-regulated gene pairs and co-expressed gene pairs. F) Validation by comparing Pearson's correlation of co-expressed and co-regulated gene pairs from Recount2 Brain in the independent GTEx and ROSMAP data.

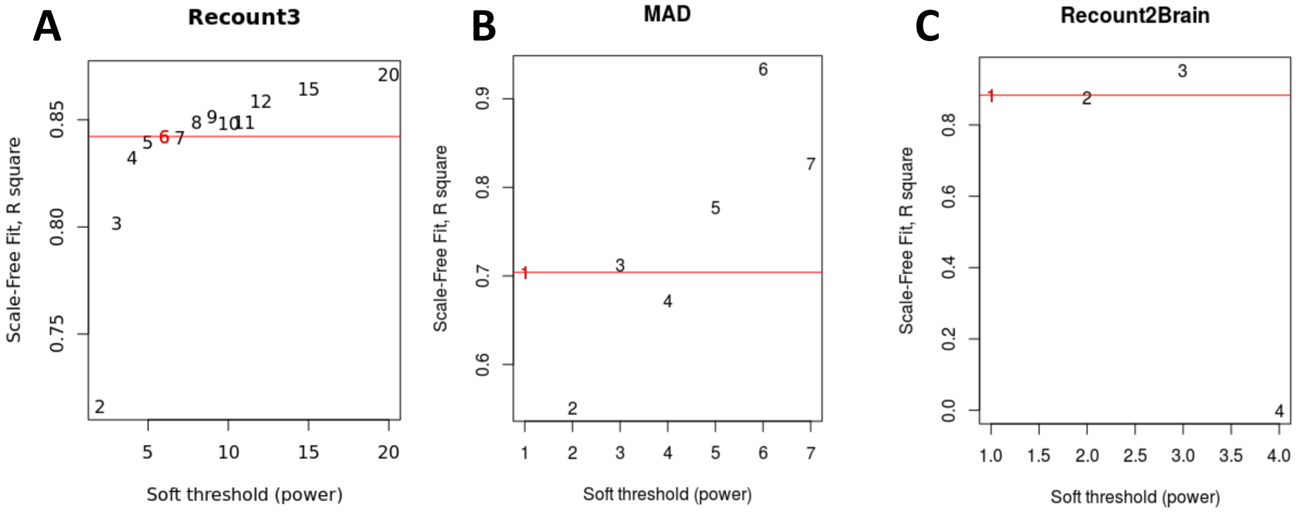


Supplementary Figure 5. The soft threshold (FDR) for scale free network. A) Recount3. B) MAD. C) Recount2 Brain

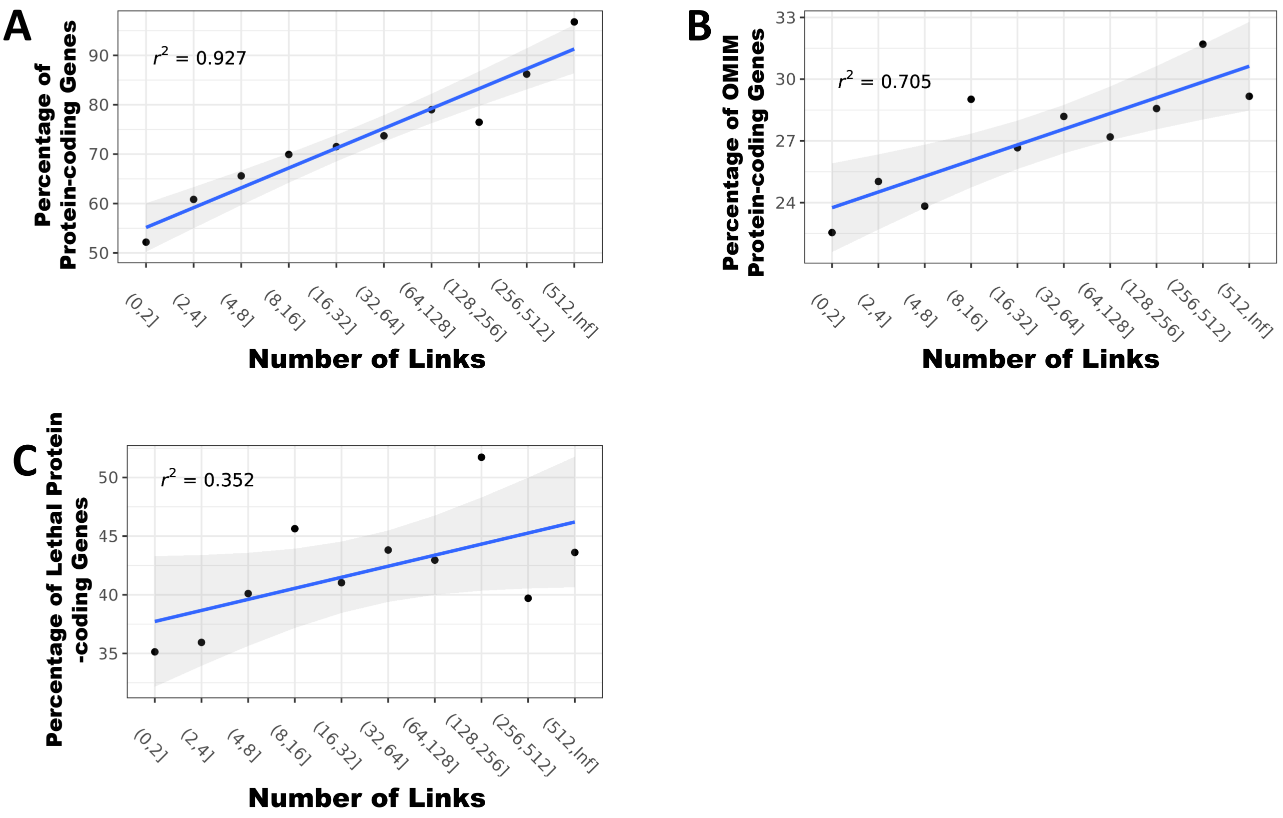


Supplementary Figure 6. Hub genes in co-regulation network are more likely to be disease-related genes A) The node degree of genes is positively related with the proportion of protein coding genes. B) The node degree of genes is positively related with the proportion of OMIM genes in protein coding genes. C) The node degree of genes is positively related with the proportion of lethal genes in protein coding genes.

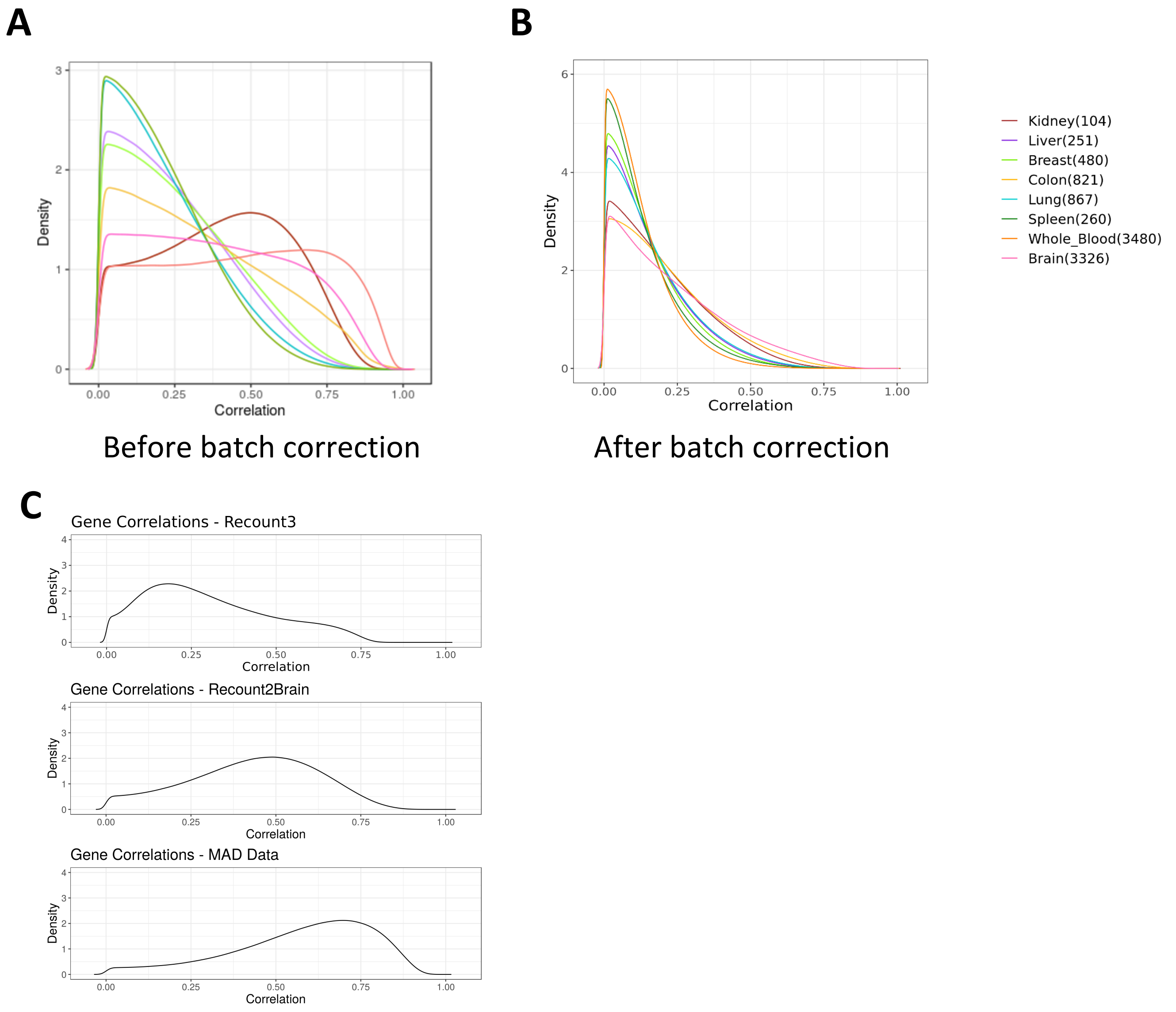


Supplementary Figure 7. Batch effect in combined data sets. A) The distribution of gene pairs' Pearson's correlations in GTEx data before batch correction. B) The distribution of gene pairs' Pearson's correlations in GTEx data after batch correction. C) The distribution of gene pairs' Pearson's correlations in Recount3, Recount2 Brain, and MAD.

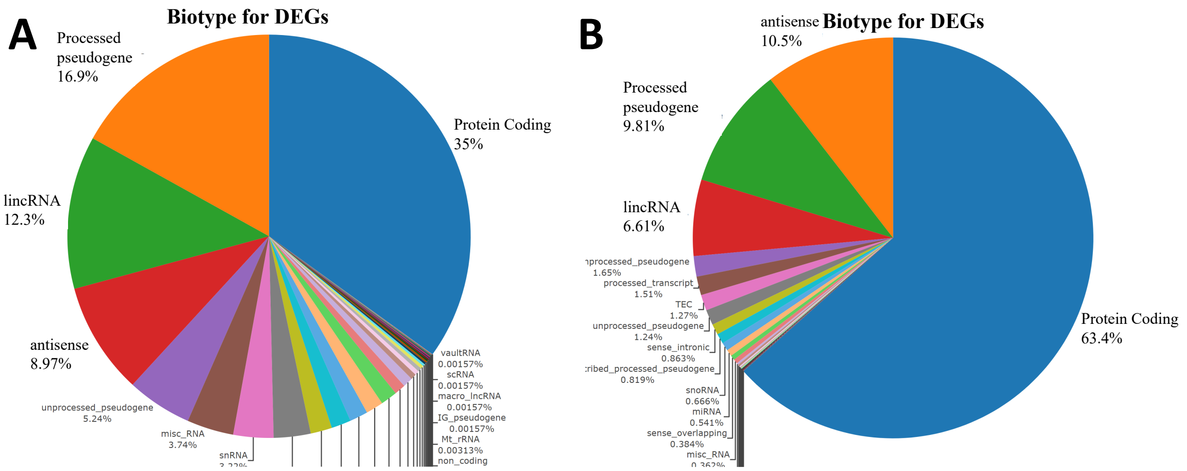


All 63,856 genes

Remaining 27,342 genes

Supplementary Figure 8: More than 27,000 genes are DEGs in more than 5% contrasts; the majority of these are protein coding genes. To save computing power, we removed genes that are DEGs in less than 5% contrasts, and we compared the biotypes of genes before (A) and after (B) removing those DEGs.

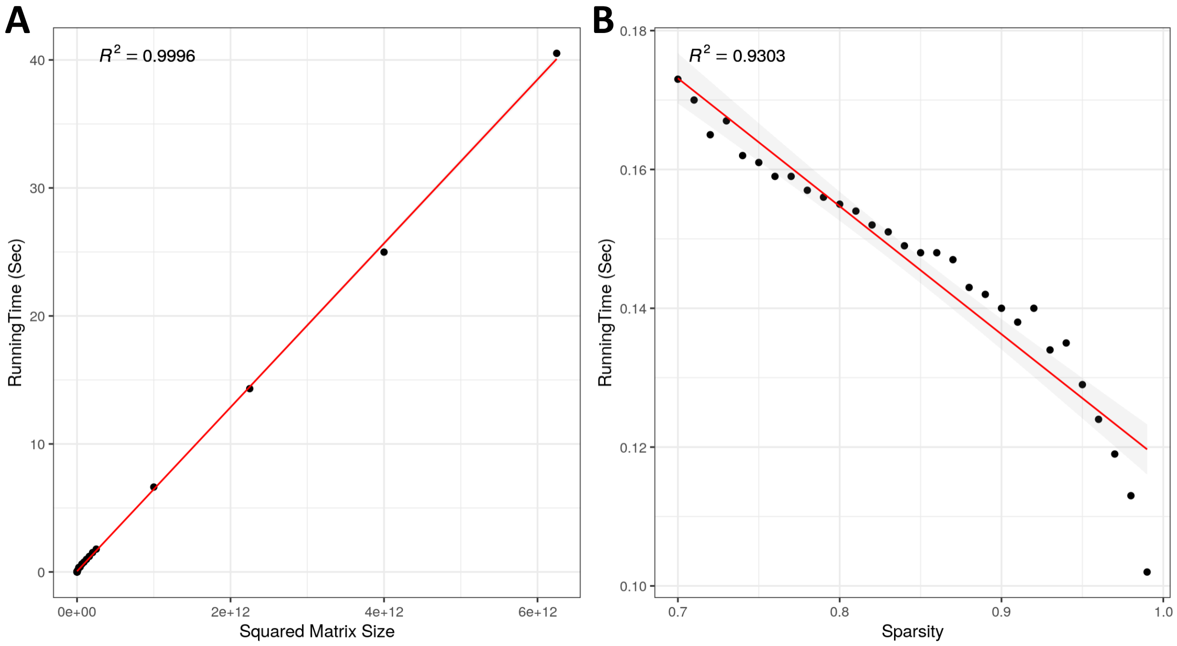


Supplementary Figure 9. The runtime of generating a random binary DEG matrix. A) The runtime is completely linearly related with the squared matrix size. B) The runtime decreased with increased matrix sparsity.

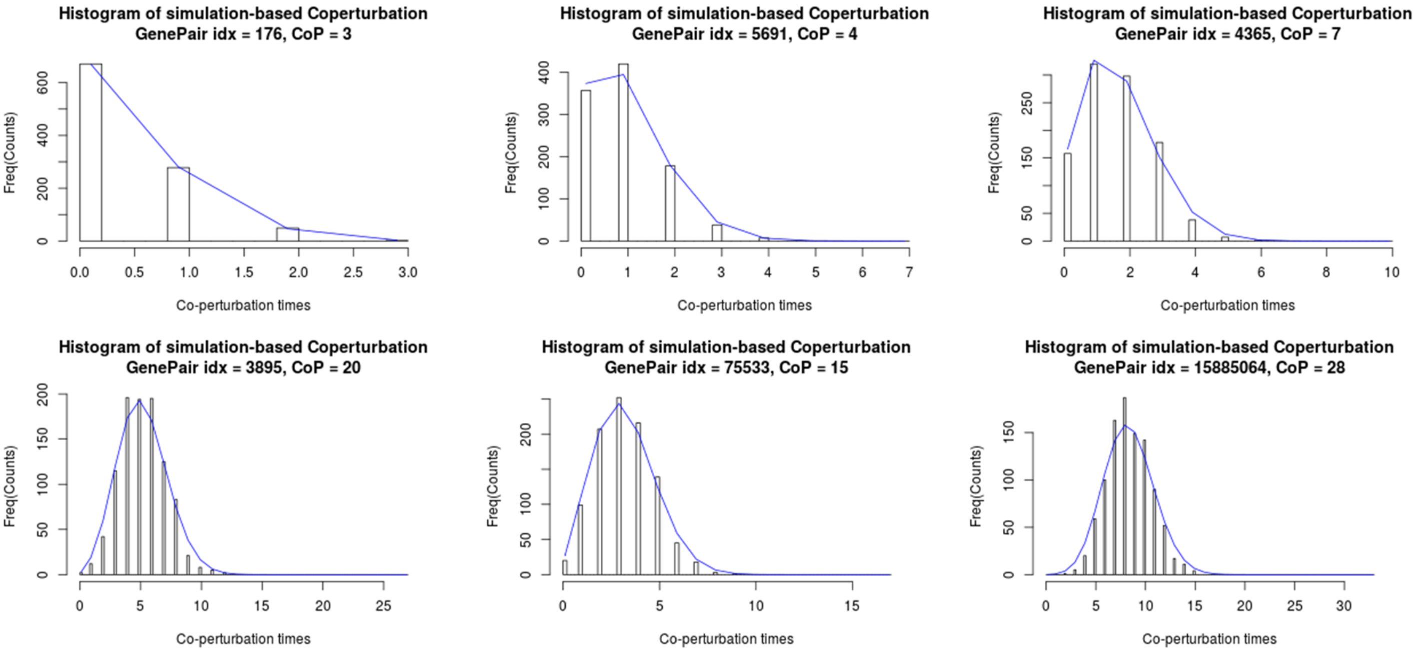


Supplementary Figure 10: Six examples that the beta-binomial fitting performance is generally good. The histograms are the simulation-based co-regulation results under random cases. The blue curves are the beta-binomial fitting curves.

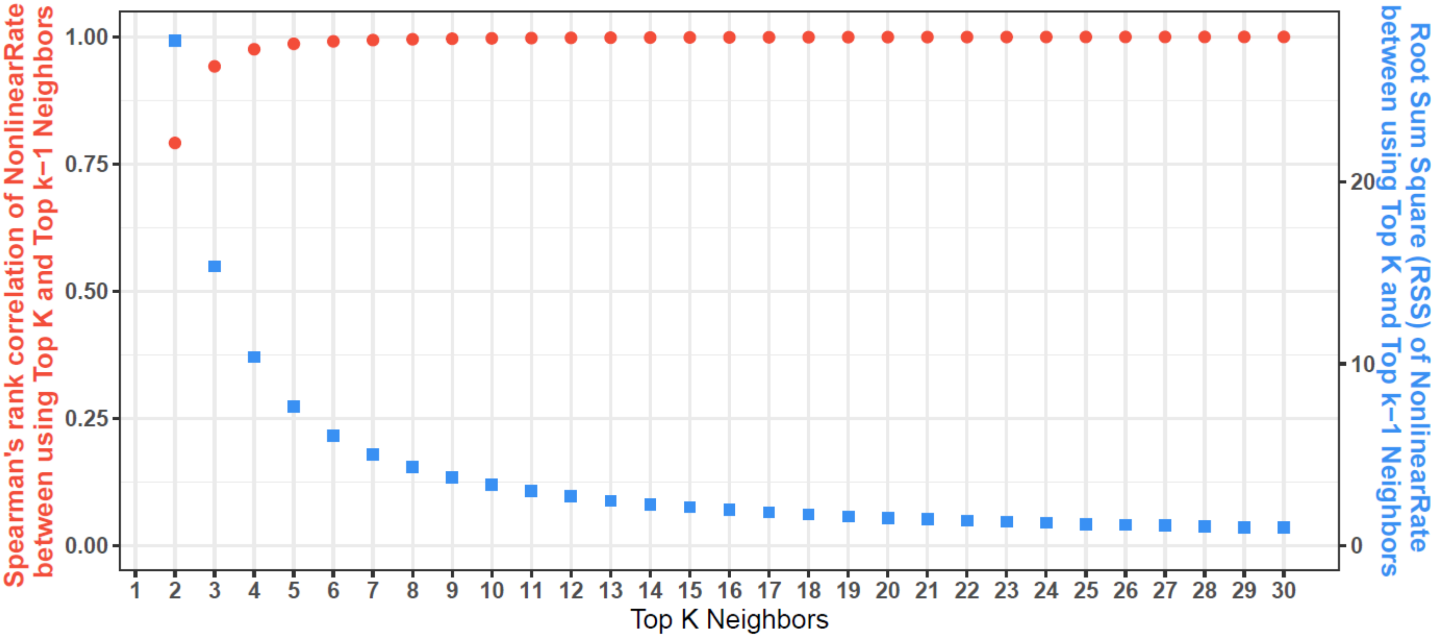


Supplementary Figure 11: The top K neighbor cutoff for determining linear and non-linear genes. The red curve shows the Spearman's correlation of gene rank between using top K and top K-1 neighbors. The blue curve shows the root sum square (RSS) of non-linear score between using top K and top K-1 neighbors.
