## Supplementary figures and images for "CoRegNet: Unraveling Gene Co-regulation Networks from Public RNA-Seq Repositories Using a Beta-Binomial Statistical Model"

### Sup Animation1

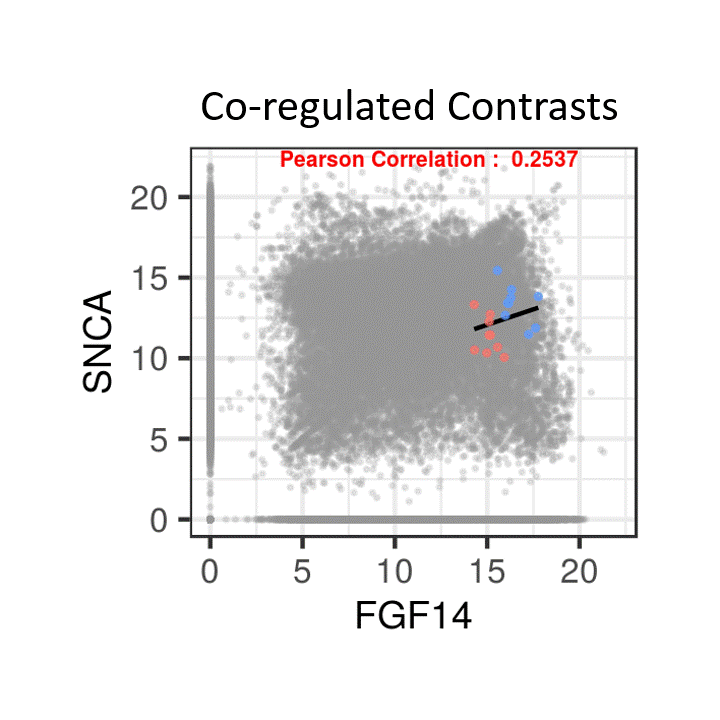
