## Supplemental Methods for "CoRegNet: Unraveling Gene Co-regulation Networks from Public RNA-Seq Repositories Using a Beta-Binomial Statistical Model"

**Supplementary Methods:**

All computational analysis was conducted in the R language (version 3.5.2 - 4.1.0).

**Data collection and processing:**

We obtained 3 datasets from 3 public databases including Recount3 (https://rna.recount.bio/), Recount2 Brain data (recount-brain project, Ramzara et al., bioRxiv, 2019), and MAD (https://doi.org/10.7303/syn2580853).

We collected 4,473 human experiments with a sample size between 6 and 40 from the Recount3 portal. We successfully extracted group label information in 3,293 experiments. For the remaining experiments, we used the DASC tool to identify hidden labels in 458 experiments. According to the label information, we found 19,681 total contrasts of which 18,997 were from manual labels and 684 were from DASC labels. These studies contained 56,498 total samples. We used DESeq2 to identify genes with fold changes greater than 1.5 and FDRs smaller than 0.01 as DEGs. Finally, we collected 12,082 contrasts with DEG numbers between 2 and 3,000, and 27,342 genes which have been identified as DEGs in at least 5% of contrasts, as our input data. Most remaining genes are protein coding genes (Sup Figure 8).

We used a similar procedure to process data from Reount2 Brain and MAD. The only difference is that genes with fold changes greater than 1.2 and FDRs smaller than 0.01 were identified as DEGs. We collected 93 contrasts from 78 experiments from Recount2 Brain, which contains ~1,300 samples and 15,000 DEGs. We also collected 138 contrasts from 70 experiments in MAD, which contain ~1,000 samples and 10,000 DEGs.

**Co-regulation matrix generation:**

Initially, we created a binary DEG matrix, where each row corresponds to a specific gene and each column represents an individual experiment. If a gene qualified as a DEG, the corresponding matrix entry was marked as 1, otherwise it was set to 0. Subsequently, we multiplied the DEG matrix by its transposed matrix to generate the gene-gene co-regulation matrix. In this co-regulation matrix, both rows and columns represent genes, and each matrix entry denotes the number of experiments in which the two corresponding genes co-regulated.

**Sampling-based co-regulation matrix generation:**

Based on the binary DEG matrix, we calculated the row and column sums. We then employed a greedy search algorithm to generate 1,000 random DEG matrices, each possessing the same row and column sums as the original binary matrix. We initiated by assigning 1s in the row or column with the highest degree of freedom, and subsequently updated this information across all rows and columns. This process was iteratively conducted until the entire matrix was populated. If at any point we encountered difficulties in assigning 1s, we discarded the entire sampled matrix and initiated a fresh iteration. For every sampled random binary DEG matrix, we performed matrix multiplication to obtain the corresponding sampling-based co-regulation matrix.

**Runtime test**

In order to pinpoint the relationship between runtime, matrix size, and matrix sparsity when generating the random binary matrix under the given row sum and column sum, we made the sparsity 0.95 (the real sparsity is usually much higher), made the column dimension 50, increased the row dimension gradually from 10 to 50,000, and recorded the runtime it took to generate a single random binary matrix. Then, we made both the row and column dimensions 1,000, increased the sparsity from 0.70 to 0.99, and recorded the runtime.

Due to the high level of sparsity in the DEG matrix, the runtime is completely linearly aligned with the squared DEG matrix size, which is the number of genes multiplied by the number of experiments (Sup Figure 9A). When the matrix size is fixed, the runtime decreases with sparsity (Sup Figure 9B). Thus, the runtime for even a large binary matrix with about 10,000 rows, 30,000 columns, and 0.99 sparsity is ≤ 1 minute.

**Beta-binomial fitting:**

Upon generating 1,000 random co-regulation matrices, we extracted the co-regulation occurrence for each gene pair from the 1,000 samplings. We applied the VGAM(34) to fit a beta-binomial distribution for every gene pair's co-regulation distribution. The accuracy of the fitting was satisfactory (Sup Figure 10), as demonstrated in Supplementary Figure 9. For each gene pair, the actual co-regulation frequency in the true co-regulation matrix was used as a threshold for the corresponding beta-binomial fitting curves. Subsequently, we calculated the p-values and the FDR. In the final co-regulation network, we connected gene pairs that exhibited significant FDRs.

**Compare Pearson’s correlation between linear and non-linear genes**

For each gene in the Recount3 co-regulation network, we ordered its neighbor by the FDR score and choose the top 20 neighbors of it to calculate the mean FDR and mean non-linear score (If the gene has less than 20 neighbors, we used all of its neighbors). As the result suggests that considering the top 20 neighbors provides a sufficiently representative measure of each gene's non-linearity (Sup Figure 11). For genes with mean non-linear score less than 0.3 and mean FDR less than ${10}^{-10}$ are identified as linear genes. For genes with mean non-linear score greater than 0.7 and mean FDR less than ${10}^{-10}$ are identified as non-linear genes. We then selected these genes along with their top 20 neighboring genes to form gene pairs. Subsequently, we evaluated the Pearson's correlation of these gene pairs across all tissue types in the GTEx dataset excluding *FrontalCortex(BA9)*, *FallopianTube*, *Cervix-Endocervix* and *Cervix-Ectocervix*.

**Ontology Tree of GO and HPO**

We constructed GO term tree but limited it to the category of 'biological process.' We then restructured the terms to ensure each node had a single parent. In cases where a node had multiple parents, it was eventually linked to the parent yielding the highest similarity score.

$$Similarity score= \frac{node\cap parent}{node\cup parent}$$

We also eliminated nodes that lacked any connection to parent nodes and pathways that were unconnected to the root. Subsequently, we utilized the 'clusterProfiler' package(35) to assess the GO term enrichment of both linear and non-linear genes. We selected the top 100 terms that were significantly enriched by either set of genes. Any GO terms unrelated to these 200 terms were subsequently removed from the tree. The enrichment score was calculated as -log10(p.adjust), where p.adjust represents the FDR of terms following the enrichment test. The cut off is where p.adjust equals 0.01. We employed a similar method to construct the HPO tree, with one key distinction: the acquisition of gene sets for each HPO term involved two mapping steps. First, we mapped HPO to the OMIM database, transitioning from phenotype to disease. Subsequently, we mapped OMIM to genes, thus transitioning from disease to genes.

**Co-regulation heatmap**

We identified contrasts wherein a gene pair was co-regulated. Contrasts that involved an upregulated gene were colored red, while those with a downregulated gene were colored blue. The color scale was established based on the log2 fold change (log2FC) of the genes.

**Scatter plot of genes in pooled Recount3 data and in co-regulated contrasts**

RNA-seq data from Recount3 was log2 transformed and combined together without applying any batch correction. We computed the overall Pearson's correlation. Contrasts where two genes were concordantly co-regulated were extracted, and samples within the same contrast were assigned the same color. Discordantly co-regulated samples were processed in a similar manner. The color generation was performed using the 'rainbow' palette in R. However, due to the vast number of contrasts, samples from distinct contrasts may exhibit similar colors.

**Module detection:**

We employed the recursive Louvain module detection method to identify modules within the co-regulation network. Initially, we executed the Louvain clustering on the entire network, resulting in several large modules for which we conducted a GO term enrichment analysis. If a module's genes were significantly enriched in multiple GO terms, we continued to apply Louvain clustering on that module to obtain more specific sub-modules. This process was repeated until the division resulted in a sub-module without any enriched GO terms. In such cases, we reverted to the previous step and retained the module in its entirety.

**Binary heatmap of gene modules**

In the heatmap, rows correspond to contrasts and a color bar distinguishes contrasts originating from different studies via unique colors. Columns represent genes, and an accompanying color bar signifies genes from distinct modules. Genes shaded in grey were not part of any module. Within the same module, genes were arranged based on their frequency of differential expression.

**GO enrichment test**

We employed the 'clusterProfiler' package to generate the plot. For each gene module, we conducted a GO enrichment test and identified the significant GO terms with an FDR less than 0.05. For each module, we only displayed the most significant GO terms. Subsequently, we utilized a dot plot as a binary matrix where each row corresponds to a GO term and each column signifies a gene module. We clustered the modules based on their Euclidean distance.

**GO term interaction density comparison:**

The Go term density was defined as Density(t) = NE(t)/NP(t), where NE(t) is the number of edges between genes assigned to term t, and NP(t) is the number of gene pairs in the term, which equals to $C_{n}^{2}$, where n is the number of genes in t.

**Validation by PPI**

Protein-protein interaction data was downloaded from the STRING database. For recount data, we applied Homo sapiens PPI data, and for the MAD study we applied Mus musculus PPI data to perform the validation. We matched the gene pairs from the co-regulation model and co-expression model with the PPI data and calculated the percentage of gene pairs validated by STRING in each model.

**Correlation distribution**

In the correlation figure in Recount3, we generated the background distribution with Pearson’s correlation of all gene pairs in the Recount3 data. We also calculated and compared Pearson’s correlations of gene pairs that the co-regulation model identified and gene pairs that WGCNA identified.

Since Recount3 covers gene data from different organs, we validated our results on GTExMaximum, which is the maximum Pearson’s correlation of the gene pair in all GTEx tissues. We first downloaded RNA seq data from the GTEx portal and then performed batch correction for each tissue type using the limma package. We checked the distribution of gene-gene Pearson’s correlations in each tissue type and removed four tissues (Bladder, Fallopian Tube, Cervix-Endocervix and Cervix-Ectocervix) with poor quality due to the small sample size. Finally, for each gene pair, we picked the maximum correlation score in all remaining tissues to represent its correlation. The background distribution contains all genes recognized by either the WGCNA or co-regulation model. For gene modules identified by WGCNA, we removed the largest module, which included more than 10,000 genes (which is almost half the total genes).

In the Recount2 Brain project, we validated our results by GTEx brain data because studies from this project were all specific to the brain. All GTEx data were batch corrected using the R package limma.

In MAD data, we matched the mouse genes with human genes first and then used batch corrected GTEx brain data and ROSMAP data to do the validation. ROSMAP data mainly contains gene expression data from patients with Alzheimer’s disease and dementia.

**Sub-network examples**

We chose the module examples based on linear rate, gene-gene association FDRs, and the number of genes. We chose the disease related module using an extremely high FDR cutoff and then matched these genes with the OMIM database.

We generated the heatmap using the same approach we used to generate the co-regulation heatmap. The only difference is that we did not scale the color per log2FC because the number of contrasts (number of rows in the heatmap) was too large. We generated the scatter plot the same way we generated the scatter plot of genes in pooled Recount3 data.

**Property of hub genes investigation**

We downloaded the disease gene label from the OMIM database. We designated all genes with disease phenotypes as disease-related genes. Then, we downloaded LoF scores of all the genes from the GnomAD database and defined the genes with LoF scores lower than 0.35 as genes intolerant of LoF alleles. The scale-free network was generated with an FDR cutoff of ${10}^{-6}$. We ordered the genes by their degree in the network and divided the degree into 9 groups with an increasing degree step of $2^{n}$. Then, we calculated the percentage of OMIM genes and of genes intolerant of mutations in each group. Finally, we made the scatter plot using gene degree as the x-axis and percentage of genes as the y-axis.

**Relationship between co-regulation times and significance**

We selected two gene pairs to show that higher co-regulation times do not always have higher significance. We plotted the histogram of co-regulation times based on 1,000 samplings and the 1-ecdf figure based on the sampling results. However, overall, higher co-regulation times yield higher significance, which can be seen from our comparison of the histogram of co-regulation times of all genes versus co-regulated genes.

**Soft threshold determination**

We applied different FDRs as cutoffs for significantly co-regulated gene pairs, and for each FDR, we generated the network and analyzed the relationship between the network degree and number of nodes in the network. We applied linear regression to log the transformed network degree and the transformed number of nodes and calculated the R square to represent the network’s scale-free property. We chose the largest FDR as the cutoff when the next smaller FDR gave us a smaller scale free fit R square.

**GTEx correlation histogram before and after batch correction**

We downloaded the original GTEx expression data and calculated the Pearson’s correlation as the before-batch-correction histogram. Then, we applied the limma package to remove the batch effect and performed the same calculation to get the after-batch-correction histogram.
